# 14-3-3ζ regulates adipogenesis by modulating chromatin accessibility during the early stages of adipocyte differentiation

**DOI:** 10.1101/2024.03.18.585495

**Authors:** SA Rial, Z You, A Vivoli, D Sean, Amal Al-Khoury, G Lavoie, M Civelek, A Martinez-Sanchez, PP Roux, TM Durcan, GE Lim

**Affiliations:** Department of Medicine, Université de Montréal, Montreal, QC, Canada; Cardiometabolic Axis, Centre de Recherche du Centre Hospitalier de l’Université de Montréal (CRCHUM), Montreal, QC, Canada; The Neuro’s Early Drug Discovery Unit (EDDU), McGill University, 3801 University Street, Montreal, QC, H3A 2B4, Canada; Institute for Research in Immunology and Cancer (IRIC), Université de Montréal, Montréal, Québec, Canada; Department of Pathology and Cell Biology, Faculty of Medicine, Université de Montréal, Montréal, Québec, Canada; Department of Biomedical Engineering, University of Virginia, Charlottesville, United States; Center for Public Health Genomics, University of Virginia, Charlottesville, VA 22908; Section of Cell Biology and Functional Genomics, Department of Metabolism, Digestion and Reproduction, Imperial College London, Hammersmith Hospital, London, UK

**Keywords:** Adipose tissue, adipogenesis, adipocyte progenitor cells, differentiation, scaffold proteins, 14-3-3ζ, ATAC-seq

## Abstract

We previously established the scaffold protein 14-3-3ζ as a critical regulator of adipogenesis and adiposity, but the temporal specificity of its action during adipocyte differentiation remains unclear. To decipher if 14-3-3ζ exerts its regulatory functions on mature adipocytes or on adipose precursor cells (APCs), we generated *Adipoq*14-3-3ζKO and *Pdgfra*14-3-3ζKO mouse models. Our findings revealed a pivotal role for 14-3-3ζ in APC differentiation in a sex-dependent manner, whereby male and female *Pdgfra*14-3-3ζKO mice display impaired or potentiated weight gain, respectively, as well as fat mass. To better understand how 14-3-3ζ regulates the adipogenic transcriptional program in APCs, CRISPR-Cas9 was used to generate TAP-tagged 14-3-3ζ-expressing 3T3-L1 preadipocytes. Using these cells, we examined if the 14-3-3ζ nuclear interactome is enriched with adipogenic regulators during differentiation. Regulators of chromatin remodeling, such as DNMT1 and HDAC1, were enriched in the nuclear interactome of 14-3-3ζ, and their activities were impacted upon 14-3-3ζ depletion. The interactions between 14-3-3ζ and chromatin-modifying enzymes suggested that 14-3-3ζ may control chromatin remodeling during adipogenesis, and this was confirmed by ATAC-seq, which revealed that 14-3-3ζ depletion impacted the accessibility of up to 1,244 chromatin regions corresponding in part to adipogenic genes, promoters, and enhancers during the initial stages of adipogenesis. Moreover, 14-3-3ζ-dependent chromatin accessibility was found to directly correlate with the expression of key adipogenic genes. Altogether, our study establishes 14-3-3ζ as a crucial epigenetic regulator of adipogenesis and highlights the usefulness of deciphering the nuclear 14-3-3ζ interactome to identify novel pro-adipogenic factors and pathways.

## INTRODUCTION

The expansion of adipocyte number occurs through adipocyte differentiation, or adipogenesis, and this process plays a pivotal role in the development of obesity (1–3). The differentiation of adipocyte precursor cells (APCs) is driven by the availability of energy-dense nutrients and a variety of adipogenic triggers, and adipogenesis is facilitated by complex signaling pathways and tightly regulated transcriptional networks (4, 5).

The canonical model of adipogenesis posits that hormonal and nutrient stimuli promote the sequential expression and activation of early adipogenic transcription factors (ATFs), which include CCAAT-enhancer-binding proteins-β/δ (C/EBP-β/δ), STAT5A signal transducer and activator of transcription 5A/B (STAT5A/B), Kruppel-like factor 5 (KLF5), and glucocorticoid receptor (GR) and late ATFs, such as C/EBPα and Peroxisome proliferator-activated receptor γ2 (PPARγ2) (4, 5). Although these events are known to take place during the early stages of APC differentiation, little is known about the molecular factors that tightly coordinate them in space (*i.e.*, cytosolic to nuclei translocation) and time (sequential activation of ATFs). This lack of knowledge has contributed to the difficulty in targeting adipogenesis with pharmacological approaches to treat obesity.

Molecular scaffold proteins likely play important roles in coordinating signaling pathways that control metabolism (6, 7), and of the various families of scaffolds, the importance of 14-3-3 proteins in signal transduction has become apparent (6, 7). Through recognition of specific phosphorylated serine or threonine motifs (RSXp<u>S/T</u>XP and RXXXp<u>S/T</u>XP), all seven mammalian 14-3-3 protein isoforms interact with a broad variety of enzymes, transcription factors, and transporters. Thus, 14-3-3 proteins can regulate diverse cellular processes, such as cell cycle progression (8, 9), apoptosis (8, 10), secretion, and metabolism (7, 9, 11–18).

Despite a high degree of sequence-homology, 14-3-3 family members can perform isoform-specific biological functions, and our group has identified critical roles of the 14-3-3ζ isoform in the regulation of whole-body adiposity, adipogenesis, adipocyte function, and glucose (7, 9, 12–17). In the context of adipocyte differentiation *in vitro*, silencing of *Ywhaz* (the gene coding for 14-3-3ζ) blocked the differentiation of 3T3-L1 pre-adipocytes (9, 14–16). We also found that systemic deletion of 14-3-3ζ significantly reduced visceral adiposity and promoted glucose intolerance and insulin resistance (9). Postnatal depletion of 14-3-3ζ in mature adipocytes was found to reduce adipose tissue *Pparg2* mRNA expression and impair the lipolytic response of adipose tissue (16). In contrast, whole-body transgenic overexpression of 14-3-3ζ potentiated age-dependent and high-fat diet-induced expansion of adipose tissue (9). Taken together, these findings demonstrate important roles of 14-3-3ζ in adipocyte development and function.

With the ability of 14-3-3 proteins to interact with a diverse array of phosphorylated proteins, we have discovered that the interactome of 14-3-3ζ changes in response to physiological and pathophysiological stimuli like adipocyte differentiation or obesity, respectively. We first used mouse embryonic fibroblasts derived from transgenic mice expressing a (TAP) epitope-tagged 14-3-3ζ molecule combined with affinity proteomics to discover that the interactome of TAP-14-3-3ζ is enriched with RNA splicing factors following the induction of adipocyte differentiation (14). Recently, we have determined that the interactome of 14-3-3ζ in adipose tissue is also sensitive to high-fat diet-induced obesity (15). Altogether, these findings demonstrate the usefulness of determining the 14-3-3ζ interactome in identifying novel regulators of adipocyte differentiation or expansion of adipose tissue mass.

A major limitation of whole-body deletion or overexpression 14-3-3ζ mouse models is the inability to distinguish the individual contributions of specific cell types, and with our findings that 14-3-3ζ may regulate adiposity (9, 16), whether 14-3-3ζ primarily influences mature adipocytes or APCs, let alone other cell types, is unclear. Moreover, the recent discovery of adipocyte and APC heterogeneity further adds to the complexity in understanding the roles of specific proteins in adipose tissue niches (19–24). Indeed, adipocytes derived from APCs, expressing Platelet-derived growth factor receptors (PDGFRs)-α and/or β,SCA1, and additional markers, represent unique sub-populations that can be influenced by age, anatomical localization and nutritional contexts (19, 20, 22–25). For example, subsets of PDGFRα+ cells with high or low expression of CD9 were found to be committed to pro-fibrotic and adipogenic cells, respectively (23). Also, CD24+ progenitors, and not CD24^-^, were characterized as having high adipogenic potential (25). The concept of heterogeneity is not restricted to rodents, as spatial technologies have revealed heterogeneity within human adipose tissues (20). Although systemic 14-3-3ζ deletion and over-expression had opposing effects on adiposity, it is not clear whether 14-3-3ζ is differentially expressed in APCs or mature adipocytes to account for the differences in fat mass.

Herein, we sought to examine the impact of deleting 14-3-3ζ in *Adipoq*+ mature adipocytes (*Adipoq*14-3-3ζKO) and in *Pdgfra*+ APCs (*Pdgfra*14-3-3ζKO) on murine adiposity. While no differences in body weights were found in *Adipoq*14-3-3ζKO mice of both sexes, *Pdgfra*14-3-3ζKO mice exhibited a sexual dimorphic effect on body weight, whereby male and female mice exhibited moderate reductions or marked increases in body weights, respectively. Moreover, enhanced adiposity and fat mass were observed in female mice. To further define the processes regulated by 14-3-3ζ in the differentiation of APCs, CRISPR-Cas9 genome editing was used to generate 3T3-L1 pre-adipocytes that express a TAP-tagged 14-3-3ζ molecule to permit the identification of the nuclear interactome of 14-3-3ζ during the early stages of adipogenesis. Chromatin remodeling was among the highest enriched biological functions, which led to the use of Assay for Transposase-Accessible Chromatin with sequencing (ATAC-seq) to assess how 14-3-3ζ influences chromatin accessibility. We found that the expression of critical adipogenic genes, such as *Fabp4*, *Adig*, *Retn*, and *Fam83a*, are correlated with 14-3-3ζ-regulated chromatin accessibility. Taken together, our study highlights the importance of 14-3-3ζ during the early stages of adipogenesis and a novel role by which it regulates adipocyte differentiation.

## RESULTS

### Adipose expression of *Ywhaz* correlates with body fat mass and insulin resistance in mice

With the finding that extreme differences in *Ywhaz* expression are associated with opposite effects on adiposity (9), we first wanted to determine if natural variations in *Ywhaz* mRNA expression are similarly correlated with adiposity and other metabolic parameters. Thus, the Hybrid Mouse Diversity Panel (HMDP) was used to explore the association of *Ywhaz* mRNA expression in perigonadal adipose tissue samples with metabolic traits. The HMDP is composed of one hundred inbred strains of male and female mice that were fed a high-fat and high-sugar (HF/HS) diet for 8 weeks and assessed for metabolic parameters such as body weight, lean and fat masses, fasting glycemia, and insulinemia (26). A significantly positive correlation between *Ywhaz* expression and insulin resistance, as measured by HOMA-IR, was detected in male and female mice after HF/HS feeding (bicor > 0.4, p < 10^-9^, **Fig.1 A and E1**). Percent fat mass before and after HF/HS diet feeding (**Fig.1 B, C, F and G**) in males (**Fig.1 B,C**) and females (**Fig.1 F,G**) were also found to be significantly correlated with *Ywhaz* mRNA expression. Altogether, these observations establish a positive association between adipose tissue expression of *Ywhaz* and increased adiposity induced by high calorie intake.

**Figure 1:**
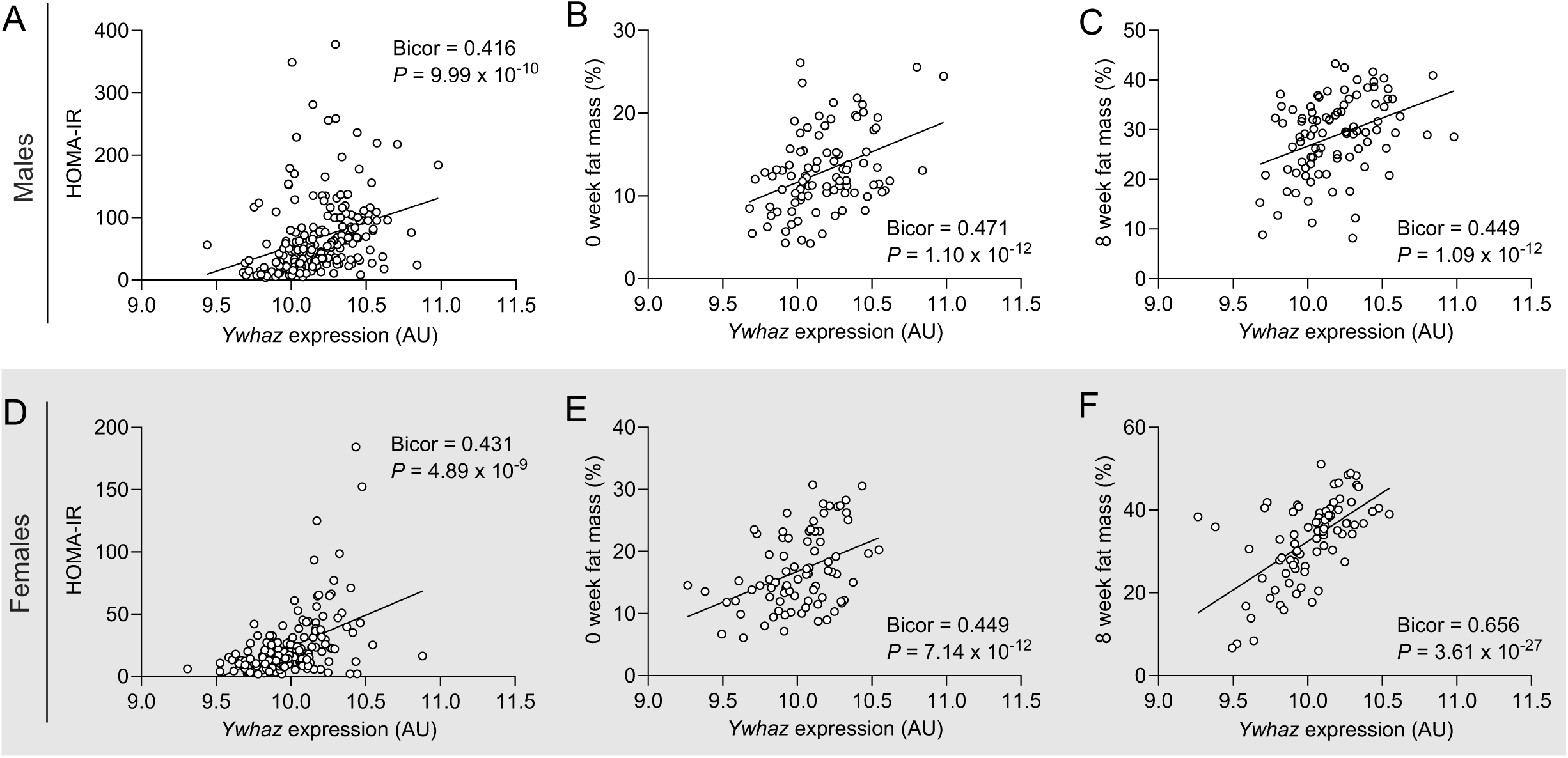
Adipose tissue expression of *Ywhaz* is positively correlated with body fat mass and insulin resistance in male and female mice. Correlation of **(A-C)** male and **(D-F)** female perigonadal adipose tissue (VAT) gene expression of *Ywhaz* with HOMA-IR (A and D) and whole body fat mass before (B and E) and after (C and F) high-fat, high-sucrose feeding, using the Hybrid Mouse Diversity Panel (HMDP) resource (26).

### Deletion of 14-3-3ζ in mature adipocytes does not impact adiposity

As a first step in determining if deletion of 14-3-3ζ in mature white adipocytes directly affects postnatal adiposity, we bred *Adipoq*^EVDR^-Cre transgenic mice (27) with *Ywhaz^flox/flox^*(13, 16), to generate mature adipocyte-specific 14-3-3ζ knockout mice (*Adipoq*14-3-3ζKO). We confirmed by quantitative PCR that *Ywhaz* expression was significantly decreased in white adipose tissues of males and female KO mice (**Fig. 2, A and K**). In males, mRNA levels of some 14-3-3 isoforms, namely *Ywhae*, *Ywhag,* and *Ywhah,* were significantly increased, but these increases were not seen in females (**Sup. Fig. 1**). Deletion of 14-3-3ζ in mature adipocytes resulted in glucose intolerance in male and female mice (**Fig. 2, C and M**); however, no differences in insulin sensitivity were detected (**Fig. 2, D and N**). Compared to WT controls, male or female *Adipoq*14-3-3ζKO mice showed no differences in body weight gain (**Fig. 2B and L**), perigonadal and inguinal adipocyte size (**Fig. 2E-G and O-Q**), or fat mass (**Fig. 2H-I and 2R-T).** Similarly, no changes in the expression of key metabolic genes, namely *Srebf1*, *Hsl*, and *Adipoq*, were detected in inguinal and perigonadal adipose tissues of both groups (**Sup. Fig. 2**). Taken together, these observations demonstrate that 14-3-3ζ in mature adipocytes is dispensable for the expansion and proliferation of adipocytes.

**Figure 2:**
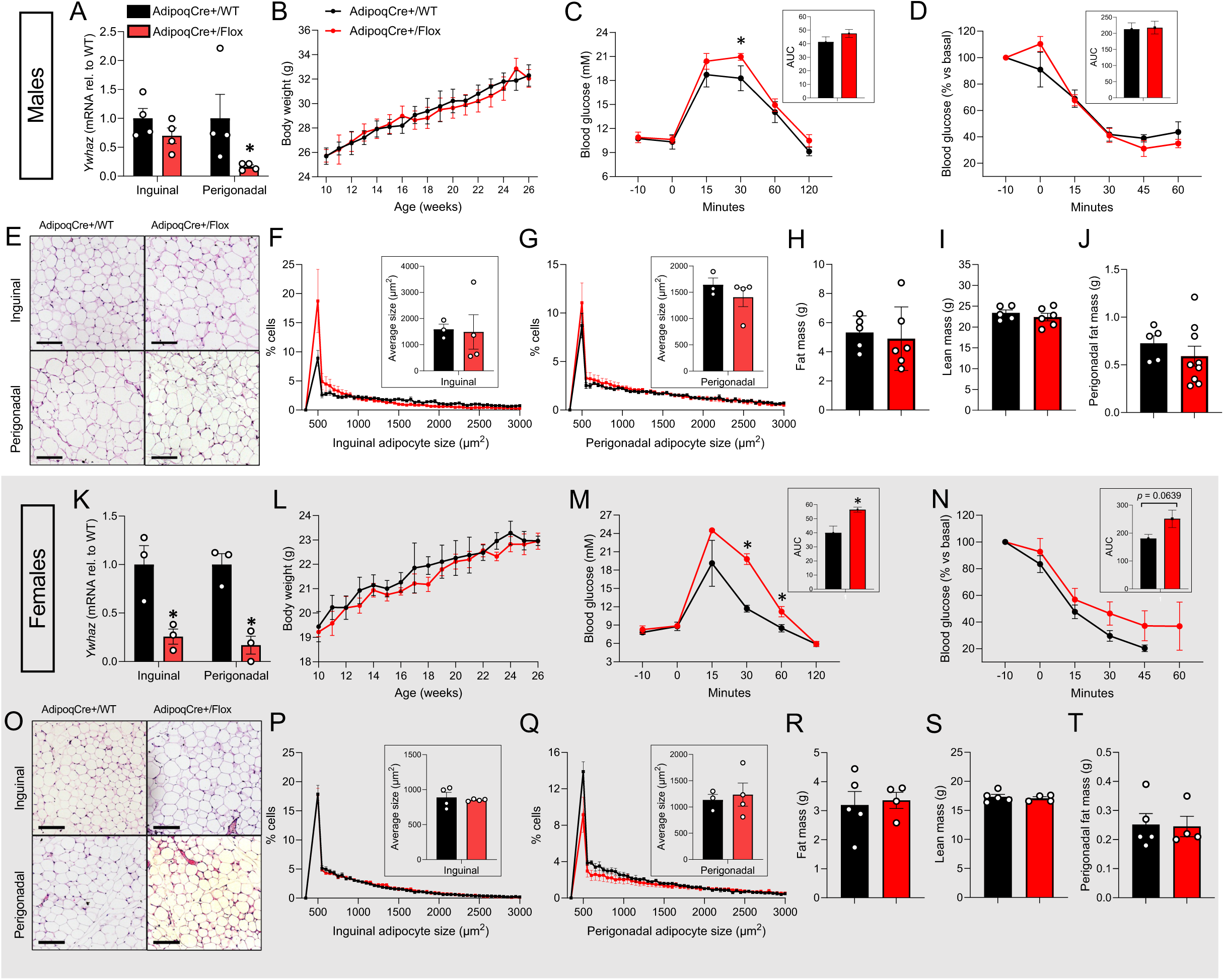
Deletion of 14-3-3ζ in mature adipocytes does not affect adiposity, but impairs glucose tolerance. **(A-J)** Male and **(K-T)** female wild type (*Adipoq*-Cre+/WT) and *Adipoq14-3-3*ζKO (*Adipoq*- Cre+/Flox) mice were generated by breeding *Adipoq*-Cre mice with either WT mice or mice harboring floxed alleles of *Ywhaz*. (A,K) Inguinal (SAT) and perigonadal (VAT) adipose tissue samples were subjected to qRT-PCR to measure *Ywhaz* mRNA levels (n=3-4 per sex per genotype). Mice (n=4 minimum) were followed for (B and L) body weight gain from 10 to 26 weeks of age, assessed for (C,M) glucose tolerance and (D,N) insulin sensitivity via ip-GTT (2g kg^-1^ B.W. D-glucose) and ip-ITT (1.0 U kg- 1 humulin® R) at 25 and 26 weeks, respectively. (E,O) H&E-stained microsections (scale bar= 200µm) from inguinal and perigonadal adipose tissues were analysed for (F,P) white adipocyte area and (G,Q) size distribution with Visiomorph™. (H-J, R-T) Body composition of mice were measured by EchoMRI^TM^ prior to sacrifice. Error bars represent S.E.M. Significant differences between wild type and *Adipoq14- 3-3*ζKO mice are indicated by \**P*<0.05 (calculated by Student’s T-test).

### Deletion of 14-3-3ζ in adipocyte progenitor yields sex-specific differences in adiposity

The lack of effect on adiposity in *Adipoq*14-3-3ζKO mice seemed contradictory with our previous finding that whole body KO of 14-3-3ζ significantly reduced visceral adiposity and adipocyte maturity (9). This suggested that 14-3-3ζ may rather play critical roles at earlier stages of adipogenesis, and more precisely in APCs. To address this hypothesis, we deleted 14-3-3ζ in APCs by breeding *Pdgfra*- Cre mice (28) with *Ywhaz^flox/flox^* to generate *Pdgfra*14-3-3ζKO mice. We confirmed by quantitative PCR that *Ywhaz* expression was significantly decreased in white adipose tissues of males and female KO mice (**Fig. 3 A and K**). As *Pdgfra*+ cells within adipose tissue are widely identified as resident adipocyte precursors (29, 30), our approach allows for the study of 14-3-3ζ in APC populations. Of note, mRNA levels of various 14-3-3 isoforms were changed in inguinal or gonadal adipose tissues in male and female mice (**Sup. Fig. 3**). Compared with WT controls, *Pdgfra*14-3-3ζKO males showed decreased body weight gain until experimental endpoint (**Fig. 3B**) and a slight tendance towards decreased adiposity (**Fig. 3H**). However, no notable effects on glucose tolerance and insulin sensitivity were observed (**Fig. 3C and D**). Deletion of 14-3-3ζ in APCs of male mice did not affect the expression of key metabolic genes (**Sup. Fig. 4A**), adipocyte size (**Fig 3. E-J**), or visceral fat mass (**Fig. 3J**).

**Figure 3:**
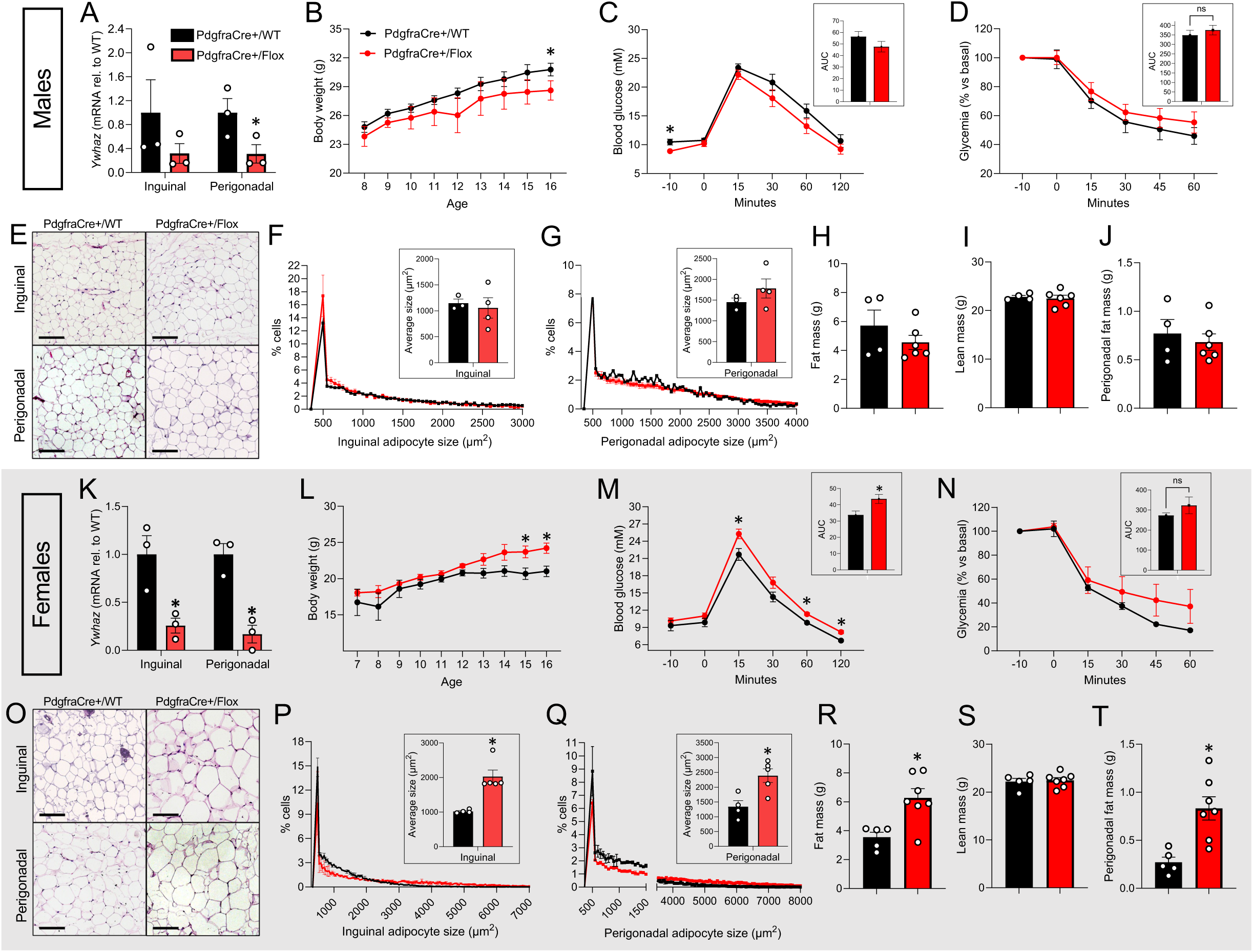
Deletion of 14-3-3ζ in adipocyte progenitor cells affects adiposity in a sex-dependent manner and impairs glucose tolerance. Male **(A-J)** and female **(K-T)** wild type (*Pdgfra*-Cre+/WT) and *Pdgfra*14-3-3ζKO (*Pdgfra*-Cre+/Flox) mice were generated by breeding *Pdgfra*-Cre mice with either WT mice or mice harboring floxed alleles of *Ywhaz*. (A,K) Inguinal (SAT) and perigonadal (VAT) adipose tissue samples were subjected to qRT-PCR to measure *Ywhaz* mRNA levels (n=3-4 per sex per genotype). (B,L) Mice (n=4 minimum) were followed for body weight gain from 10 to 26 weeks of age, assessed for (C,M) glucose tolerance and (D, N) insulin sensitivity via ip-GTT (2g kg^-1^ B.W. D-glucose) and ip-ITT (1.0 U kg-1 humulin® R) at 25 and 26 weeks, respectively. (E, O) H&E-stained microsections (scale bar= 200µm) from inguinal and perigonadal adipose tissues were analysed for (F, P) white adipocyte area and (G, Q) size distribution with Visiomorph™. (H-J, R-T) Body composition of mice were measured by EchoMRI^TM^ prior to sacrifice. Error bars represent S.E.M. Significant differences between wild type and *Pdgfra14-3-3*ζKO mice are indicated by \**P*<0.05 (calculated by Student’s T-test).

Surprisingly, *Pdgfra*14-3-3ζKO female mice exhibited significantly increased body weight by 16 weeks of age (**Fig. 3L**). Impaired glucose tolerance was observed following an intraperitoneal glucose bolus (**Fig. 3M**), and no differences in insulin sensitivity were observed between genotypes (**Fig. 3N**). Notably, the average size of inguinal and perigonadal adipocytes in female *Pdgfra*14-3-3ζKO mice were significantly increased by up to 2-fold (**Fig. 3O and R**), and indeed, larger adipocytes (>2,500µm^2^) accounted for up to 40% of total inguinal and perigonadal adipocytes in *Pdgfra*14-3-3ζKO females, while their frequency did not exceed 14% in WT females (**Sup. Fig. 5**). Consistently, whole-body fat mass was also significantly increased by 1.7-fold (**Fig. 3R**), and perigonadal WAT mass increased by 3-fold (**Fig. 3T**). No effects on lean mass were observed following 14-3-3ζ deletion in APCs (**Fig. 3S**). *Srebf1* gene expression was significantly decreased in both fat depots of *Pdgfra*14-3-3ζKO female mice, and expression of the lipolytic gene, *Hsl,* was significantly downregulated only in perigonadal fat (**Sup. Fig. 4B**).

Collectively, these observations suggest that 14-3-3ζ expression in APC contributes to body weight gain in males, while restricting adipogenesis in females. Moreover, our data demonstrate that 14-3-3ζ influences adipogenesis at the level of APCs, rather than by influencing mature adipocytes.

### 14-3-3ζ is heterogeneously expressed by mature adipocytes and APCs

The limited impact on adiposity and body weight due to 14-3-3ζ deletion in APCs of *Pdgfra*14-3- 3ζKO male mice was unexpected (**Fig. 3A-J**), as we anticipated that deletion in APCs would block adipogenesis similar to our previous findings in 3T3-L1 pre-adipocytes (9). Hence, we hypothesized that 14-3-3ζ might be heterogeneously expressed in different APCs populations, as well as in certain mature adipocyte niches. Indeed, we had previously observed heterogenous expression of 14-3-3ζ in mature adipocytes in gonadal adipose tissues of male mice (16). We took advantage of the publicly available mouse white adipose tissue atlas by Emont *et al.* (31) (**Sup. Fig. 6**), and single-nuclei RNA-Seq analysis of mature adipocyte and APC clusters revealed that a small fraction of cells expresses *Ywhaz* (**Sup. Fig. 6**). An important caveat is that it is not possible to distinguish cells not expressing *Ywhaz* from dropouts due to the inherent low reads per nuclei from single-nuclei sequencing. Mature adipocytes with detectable levels of *Ywhaz* mRNA represented nearly 15% and 30% of total mature adipocytes in males in females, respectively (**Sup. Fig. 6A-E**), and APCs with detectable levels of Ywhaz represented 17% and 19% of APCs in males in females, respectively (**Sup. Fig. 6-J**). We therefore aimed to further define the regulatory mechanisms underlying 14-3-3ζ-dependent adipogenesis.

### Generation of TAP-tagged 14-3-3ζ expressing 3T3-L1 cells to interrogate the nuclear 14-3-3ζ interactome

14-3-3 proteins are not *bona fide* transcription factors as they do not contain DNA-binding domains (32). With the ability of 14-3-3 proteins to interact with metabolic transcription factors, such as FOXO1 and the CREB coactivators CRTCs (33, 34), it seemed plausible that 14-3-3ζ may influence the differentiation of preadipocytes by binding to and nucleating adipogenic transcriptional complexes. In fact, we previously showed that 14-3-3ζ interacts with C/EBP-β during adipogenesis(9). As we have previously reported that mass spectrometry can be used to elucidate the 14-3-3ζ interactome under physiological and pathophysiological contexts (14, 15), we sought to define the nuclear 14-3-3ζ interactome during the differentiation of preadipocytes, with the goal of identifying novel adipogenic transcription factors or co-regulators that could be anchored by 14-3-3ζ.

To this end, CRISPR-Cas9 editing was used to generate TAP (FLAG-HA)-tagged 14-3-3ζ- expressing 3T3-L1 pre-adipocytes (TAP-3T3-L1, **Fig. 4A-F; Sup. Fig. 7**). This strategy was chosen to avoid artifacts associated with protein overexpression, as the TAP-14-3-3ζ molecule is expressed at endogenous levels (35, 36) (**Fig. 4G Sup.Fig. 7E**). Using non-edited (WT) 3T3-L1 cells as a comparator, we confirmed that CRISPR-Cas9-edited TAP-3T3-L1 cells maintained their adipogenic potential (**Fig. 4A-F**). This was further validated by Oil Red-O staining (**Fig. 4A and B**), triglyceride storage (**Fig. 4C**), responsiveness to lipolytic stimuli (**Fig. 4D**), and glycerol release (**Fig. 4E**). Moreover, expression and nuclear accumulation of PPARγ1/2 after induction of adipogenesis with the established cocktails of MDI and MDIR were unaltered (**Fig. 4F; Sup. Fig. 7F)** (9, 16, 37). We also confirmed that knock-in of TAP-tag did not affect the expression of the remaining 14-3-3 isoforms (**Sup. Fig. 7G**).

**Figure 4:**
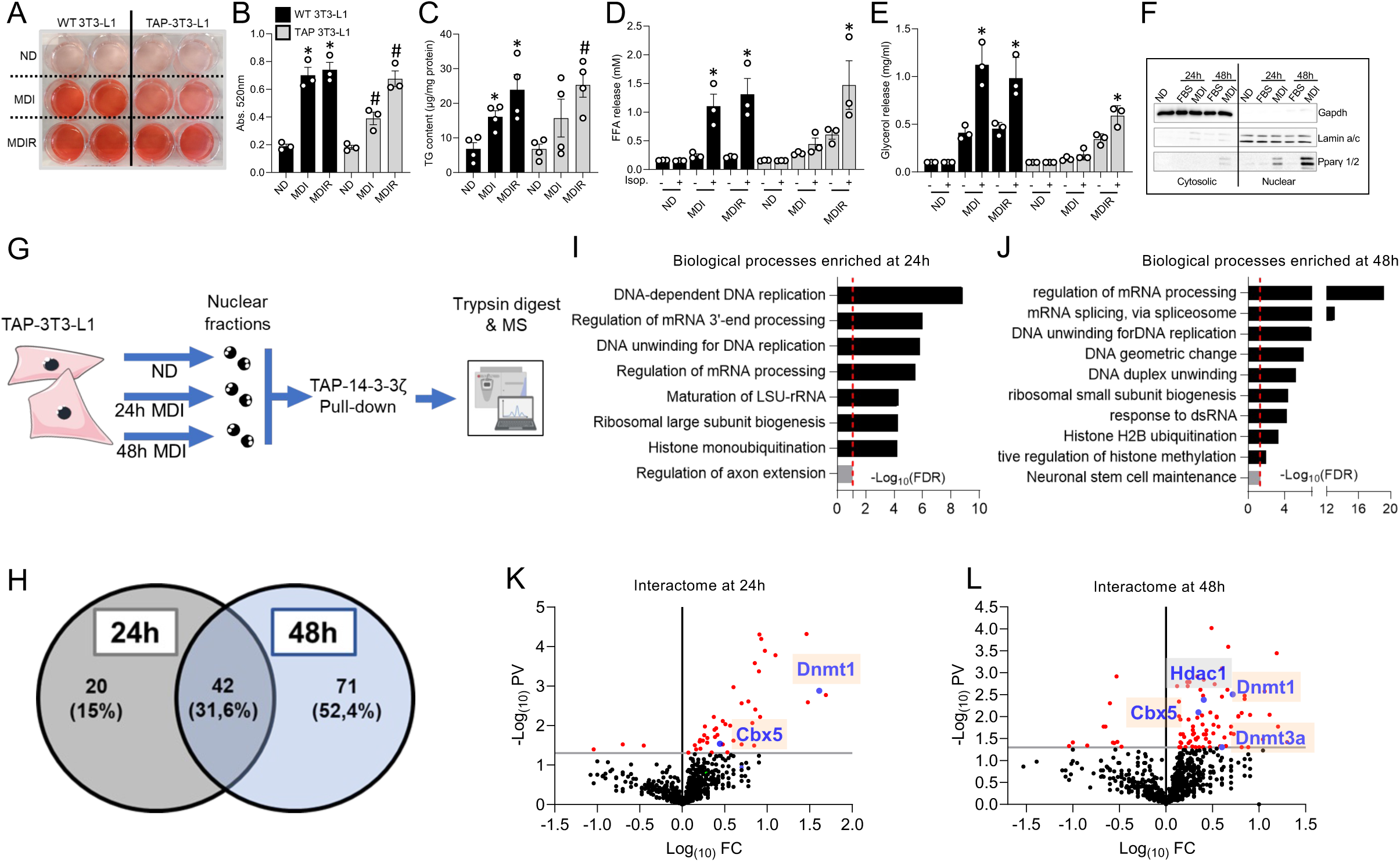
TAP-3T3-L1 cells generated by CRISPR-Cas9 allows for the elucidation of 14-3-3ζ nuclear interactome during the early stages of adipogenesis. **(A-F)** Confirmation of the ability of TAP-3T3-L1 pre-adipocytes to undergo adipogenesis following induction with MDI or MDIR by Oil Red- O (ORO) staining (A) and accompanying quantification (B), triglyceride content (C), FFA (D) and glycerol release (E) in response to isoproterenol (1µM), or, and (F) nuclear abundance of PPARγ1/2 isoforms 2 days after induction with MDI (n=3 per group per treatment). **(G)** Schematic outline of how TAP-3T3-L1 cells were used to elucidate the nuclear interactome of 14-3-3ζ during the first 24 and 48 hours of adipogenesis (n=4 for each condition). **(H)** Venn diagram showing the number of unique and overlapping proteins enriched at 24h and 48h post induction. **(I,J)** GO analysis showing the most enriched Biological Processes attributed to 14-3-3ζ interactome at 24h (I) and 48h (J) of adipogenesis. **(K and L)** Volcano plots of the decreased and enriched proteins in the 14-3-3ζ nuclear interactome. Error bars represent S.E.M. Significant differences between MDI or MDIR-induced WT or TAP-3T3-L1 cells with non-differentiating controls are indicated by \**P*<0.05 or ^#^*P*<0.05, respectively (calculated by Student’s T-test).

### Nuclear 14-3-3ζ interactors are associated with ribosomal biogenesis, RNA processing, DNA replication, and chromatin architecture during adipogenesis

Affinity proteomics was performed on nuclear TAP-14-3-3ζ complexes collected from TAP-3T3-L1 cells after 24h and 48h treatment with MDI (**Fig. 4G**). These time points were chosen to identify proteins that associate with 14-3-3ζ during the first 48h of adipogenesis, the critical, early period when signaling events necessary for murine adipogenesis occur (4, 5, 9). At 24h and 48h post-differentiation, mass spectrometry analysis revealed that the 14-3-3ζ nuclear interactome was significantly enriched with 62 and 113 proteins, respectively (FC ≥ 1.5, p ≤ 0.05; **Fig. 4H; Sup. Tables 1 and 2**). Contrary to our initial objective of discovering novel adipogenic transcription factors or regulators, the only transcription factor that we detected was C/EBP-β (**Sup.Tables 1 and 2**). To gain insights into the functional roles of identified proteins in the TAP-14-3-3ζ interactome at both timepoints, gene ontology analysis was performed. Nuclear 14-3-3ζ interacting proteins were primarily associated with histone H2B ubiquitination, DNA unwinding for replication, DNA hypermethylation and ribosome assembly (FDR ≤ 0.001, Log_10_Fold Enrichment ≥ 1.0) (**Fig. 4I and J**). We also observed proteins involved in RNA splicing and maturation, in line with our previous finding that 14-3-3ζ interactome contains RNA-splicing factors playing essential roles in adipogenesis (14).

To gain an understanding of the abovementioned biological functions in adipogenesis, we assessed how interfering in these processes would affect adipocyte differentiation (**Sup. Fig. 8A-D**). As revealed by ORO staining, Etoposide, a classic inhibitor of DNA unwinding, as well as Acivicin and Streptonigrin, two inhibitors of ribosome biogenesis and assembly (38, 39), significantly impeded 3T3- L1 differentiation when administered during the first 48h or during the full process of differentiation (**Sup. Fig. 8A and B**). Exposure of 3T3-L1 cells to Tacedinaline and Ciprofloxacin, which are inhibitors of Histone deacetylation and minichromosome maintenance proteins (MCM2-7), respectively, potentiated adipogenesis (**Sup. Fig. 8A and B**). Of note was the observation that MCM proteins were found to be among the most highly enriched proteins in the nuclear 14-3-3ζ interactome (**Sup. Table 1 and 2**). We also used siRNAs to deplete various 14-3-3ζ interactors with central roles in DNA unwinding and replication (MCM2-7) (40, 41), histone modifications (CTR9, PAF1 and LEO1) (42–44), RNA processing (NOP2) (45) and ribosome assembly (PES1, RSL1D1 and LAS1) (46, 47) (**Sup. Fig. 8C and D**). Depletion of these proteins mostly impaired adipogenesis as assessed by ORO staining (**Sup. Fig. 8C**) and *Pparg2* expression (**Sup. Fig. 8D**), except for depletion of LEO1 which significantly potentiated *Pparg2* expression. These findings support important regulatory roles of nuclear 14-3-3ζ interactors on adipogenesis.

### 14-3-3ζ regulates the activity of DNMTs, HDACs and HATs

With the detection of significantly enriched DNA methyltransferases 1 and 3a (DNMT1 and DNMT3A, respectively, Chromobox protein homolog 5 (CBX5) and Histone deacetylase 1 (HDAC1) abundance within the 14-3-3ζ nuclear interactome (**Fig. 4K and L**), this suggested that 14-3-3ζ might regulate the activity of these key epigenetic modifiers (48, 49). We therefore confirmed that silencing of *Ywhaz* in 3T3-L1 cells strikingly decreased by 4-fold the activity of DNMT enzymes (**Fig. 5B**). We also found that total HDAC activity was significantly increased by a 1.5-fold after 24h and 48h of differentiation of 3T3-L1 cells treated with siCTL (**Fig. 5C**); however, this increase was abrogated when cells were treated with si*Ywhaz* (**Fig. 5C**), suggesting a functional interaction between 14-3-3ζ and HDAC enzymes during the early stages of differentiation. Although we did not identify histone acetyltransferase (HAT) in the 14-3-3ζ interactome, we explored its dependence on *Ywhaz* expression, since it is an important modulator of histone acetylation by acting as an antagonist to HDACs. Interestingly, silencing of *Ywhaz* resulted in a significant increase of HAT activity by a 2-fold at 48h of differentiation; an increase not found in differentiating cells normally expressing *Ywhaz* (**Fig. 5D**). Overall, silencing of *Ywhaz* at early timepoints of adipogenesis blocked HDACs activity and increased HAT activity. Moreover, these results support that 14-3-3ζ, through direct or indirect interactions, balances the activity of key effectors of chromatin architecture and accessibility during the early stages of adipogenesis.

**Figure 5:**
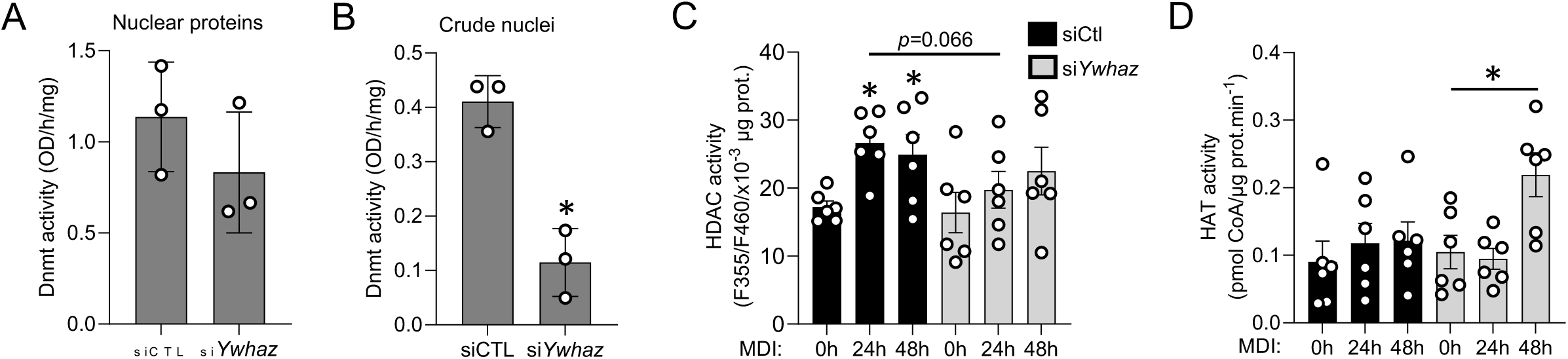
14-3-3ζ influences key effectors of chromatin accessibility during the early stages of adipogenesis. **(A,B)** Nuclear protein fractions (A) and crude nuclear fractions (B) extracted from 3T3- L1 cells transfected for 48 hours with siCTL (10 nM) or si*Ywhaz* (10 nM) were used for assessing DNMT activity, normalized by total protein (n=3 for each condition). **(C,D)** Nuclear protein fractions extracted from 3T3-L1 cells after 0, 24 or 48h of induction with MDI in presence of either siCTL or si*Ywhaz* (10 nM each) were assessed for (C) HDAC and (D) HAT activities, normalized to total protein concentration, using fluorimetric and colorimetric assays respectively. Error bars represent S.E.M. Significant differences between MDI-induced 3T3-L1 cells with non differentiating controls are indicated by \**P*<0.05 (calculated by Student’s T-test).

### 14-3-3ζ influences chromatin remodeling during adipogenesis

To explore if 14-3-3ζ regulates chromatin accessibility during adipocyte differentiation, we performed ATAC-seq on siCTL- or si*Ywhaz*-transfected 3T3-L1 cells undergoing MDI-induced differentiation (**Fig. 6A**). Compared to siCTL-transfected cells, 14-3-3ζ depletion decreased accessibility of 215, 1,244 and 943 distinct chromatin regions at 0h, 24h and 48h post-MDI induction, respectively (fold change ≤ 0.50 and *p. adj.* ≤ 0.05) (**Fig. 6B**). Among these chromatin regions with decreased accessibility, 34 and 353 overlapped with either the three timepoints or the 24h and 48h timepoints (**Fig. 6B**). Conversely, increased accessibility of 138, 1,756 and 821chromatin regions (fold change ≥ 1.50 and *p. adj.* ≤ 0.05) was seen at the three timepoints, respectively, with 22 and 461 overlapping with the three timepoints and the 24h and 48h timepoints, respectively (**Fig. 6C**). These results indicate that depletion of 14-3-3ζ during the early stages of adipogenesis reshapes the landscape of differentially accessible chromatin regions.

**Figure 6:**
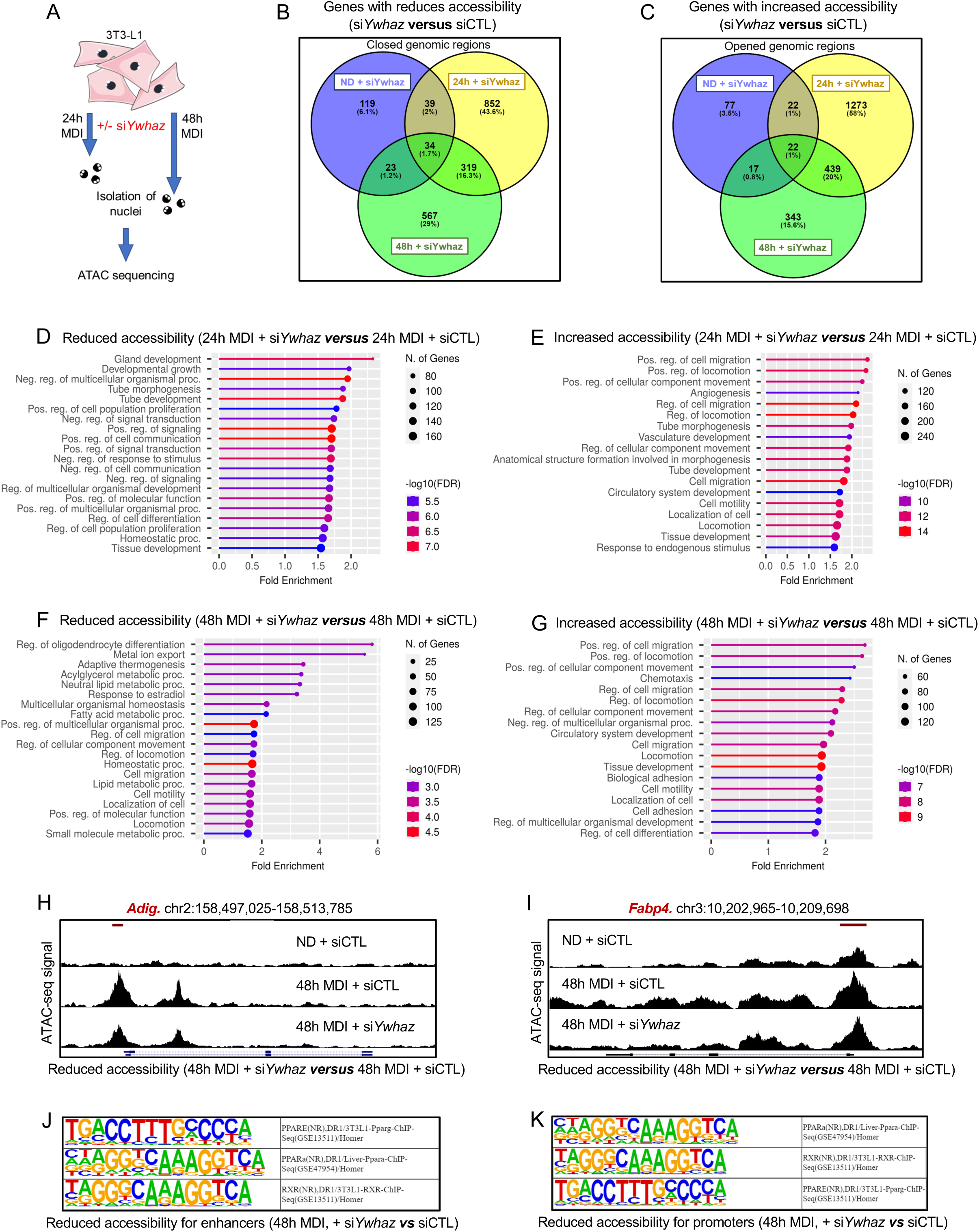
14-3-3ζ is involved in chromatin remodeling and accessibility during adipogenesis. **(A)** Schematic overview of the use of 3T3-L1 pre-adipocytes for ATAC-seq, 24h and 48h post-differentiation with MDI in the presence of siCTL or si*Ywhaz* (10nM each). **(B,C)** Venn diagrams showing overlap of (B) down-regulated or (C) differentially accessible regions after si*Ywhaz* treatment at 0, 24 and 48h post-MDI induction. **(D-G)** GO enrichment analysis with significance set to FDR ≤ 0.001 and limited to the top 20 most significant categories showing the top 20 most enriched Biological Processes gene sets corresponding to chromatin regions with reduced (D,F) and increased (E,G) accessibility after 24h (D,E) and 48h (F,G) of differentiation in presence of si*Ywhaz*. **(H,I)** Coverage tracks of ATAC-seq signals at the *Fabp4* and *Adig* genes. **(J,K)** Enhancers (J) and promoters (K) motif enrichment analysis at 48h post-MDI induction in presence of si*Ywhaz*.

Using the GREAT (v.4.0.4) region-to-genes association online tool (50, 51), we further associated these differentially accessible chromatin regions with genes and annotated these genes by Gene Ontology (- log_10_FDR ≥ 3) (**Fig. 6D-G**). Genes associated with “development” and “regulation of cell differentiation” were among those whose accessibility were decreased at 24h post MDI and after *Ywhaz* silencing (**Fig. 6D**). After 48h of differentiation, depletion of *Ywhaz* decreased accessibility of genes associated with adipocyte processes, including adaptive thermogenesis, neutral lipid metabolic processes, and lipid metabolic processes (**Fig. 6F**). Genes with increased accessibility at both timepoints in presence of si*Ywhaz* were not related to adipocyte function or development (**Fig. 6E and G**). As an example, coverage tracks of ATAC-seq signals shows significantly reduced accessibility, by depletion of *Ywhaz* 48h post-differentiation induction, of two genes involved in adipogenesis or adipocyte function, namely *Adig* (promoter region), which encodes Adipogenin (52), and *Fabp4* (exon 1), which encodes Fatty acid binding protein 4 (53, 54),, when 14-3-3ζ was depleted (**Fig. 6H and I**).

Further motif enrichment analysis revealed that when compared to siCTL-treated cells, depletion of 14-3-3ζ decreased accessibility of 47 enhancers (overlapping with H3K27ac marks and lacking H3K4me3 marks) and 16 promoters (overlapping with H3K4me3 marks), and increased accessibility of 51 enhancers and 3 promoters 24h post-MDI induction. 48h post-MDI induction depletion of 14-3-3ζ decreased accessibility of 3028 enhancers and 8 promoters.14-3-3ζ depletion also significantly (-log_10_p ≥ 5) decreased accessibility of promoters and enhancers of Peroxisome proliferator-activated receptor alpha (*Ppara*), Retinoic X receptor (*Rxr*) and Peroxisome proliferator-activated receptor elements (*PPARE*) (**Fig. 6J and K**), at 48h of differentiation of 3T3-L1 cells.

### 14-3-3ζ enables both accessibility and expression of adipogenic genes at early stages of adipogenesis

To assess how changes in chromatin accessibility correlate with changes in gene expression in the context of 14-3-3ζ-regulated adipogenesis, we integrated our ATAC-seq results with our previously published RNA-seq dataset (GSE60745, **Fig. 7A**) derived from 3T3-L1 cells treated for 0h, 24h or 48h with MDI in presence of either siCTL or si*Ywhaz* (9). These conditions correspond to what was used in our proteomic and ATAC-seq studies (**Fig. 4G and 6A**). Our integrated analysis of ATAC-seq and RNA-seq did not reveal correlations between chromatin accessibility and changes in genes expression due to 14-3-3ζ silencing at 24h of differentiation (**Fig. 7B**). However, following 48h of differentiation, silencing of 14-3-3ζ resulted in a positive and significant correlations (r>0.4, p≤0.05) between differentially accessible promoters and differentially expressed genes (**Fig. 7C**). Interestingly, *Fabp4* was found to be among the genes negatively regulated for both promoter accessibility and RNA expression at 48h of differentiation (**Fig. 7C**). Moreover, *Retn* and *Fam83a*, which respectively encode Resistin and Family with sequence similarity 83 A (Fam83A) proteins, were also among the genes with decreased promoter accessibility and expression at the same timepoint (**Fig. 7C**). Both *Retn* and *Fam83a* have previously been established as strictly required for adipogenesis (55, 56). Taken together, these findings confirmed that at early stages of adipogenesis, 14-3-3ζ is a permissive factor that promotes a landscape of chromatin accessibility that enables the expression of adipogenic genes.

**Figure 7:**
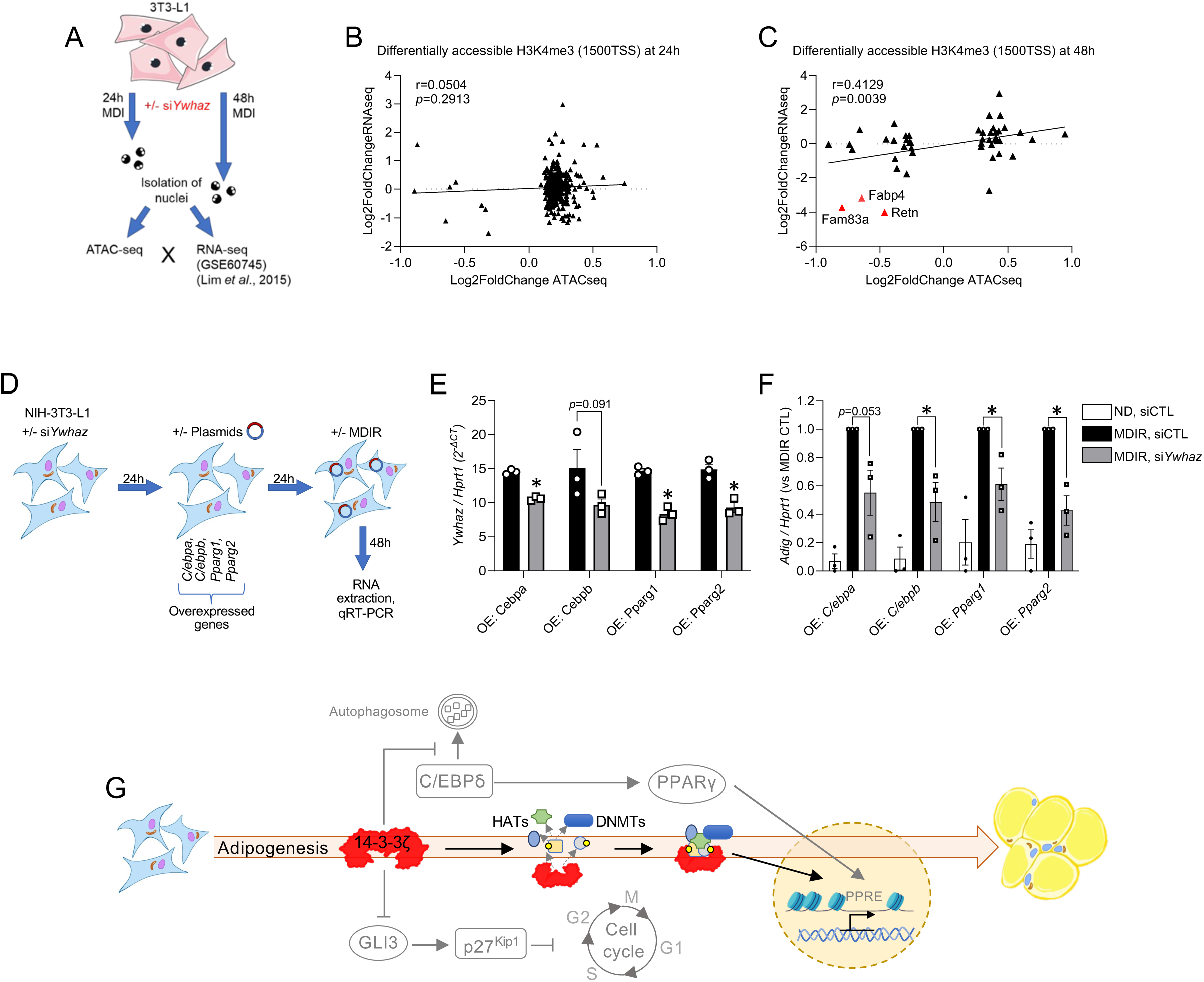
14-3-3ζ enables accessibility and the transcription of genes involved in adipocyte function and maturation. **(A)** Over-laying of ATAC-seq results with previous RNA-seq results (GSE60745) to correlate differentially accessible promoter regions with fold-changes in expression of corresponding genes, **(B)** 24h and **(C)** 48h post-MDI induction in presence of si*Ywhaz*. **(D)** Experimental outline to assess requirement of 14-3-3ζ for active ATF-dependent expression of adipogenic genes. **(E)** *Ywhaz* and **(F)** *Adig* genes mRNA expression levels in NIH-3T3 cells successively transfected with si*Ywhaz* (60 nM) and plasmids for *Cebpa, Cebpb, Pparg1,* and *Pparg2* prior to treatment with MDIR for 48h. **(G)** Graphical illustration of the new proposed mechanism (bright colors) on how 14-3-3ζ regulates adipogenesis via chromatin remodeling in integration with previously found mechanisms (dim colors) (9).

In order to further confirm that 14-3-3ζ regulates the accessibility of chromatin regions of pro-adipogenic genes, we explored if over-expression of key adipogenic ATFs could rescue the effect of 14- 3-3ζ depletion. We hypothesized that excessive over-expression of an ATF would be unable to promote differentiation if essential chromatin regions remain inaccessible. To this end, we used NIH-3T3 fibroblasts that normally lack the ability to differentiate into adipocytes. However, when ectopic ATFs such as C/EBP-β, C/EBP-α, or PPARγ2 are over-expressed, they acquire the capacity to undergo spontaneous, MDI-induced, or/and PPARγ agonism-dependent adipogenesis (57–59).

Accordingly, NIH-3T3 cells were sequentially transfected with or without si*Ywhaz* and plasmids encoding C/EBPα, C/EBPβ, PPARγ1 or PPARγ2, followed by MDIR to induce differentiation (**Fig. 7D-F**). Up to 10-fold induction of *Adig* gene expression was detected in MDIR-treated NIH-3T3 cells overexpressing each ATF (**Fig. 7F**); however, when 14-3-3ζ was depleted, *Adig* expression was significantly blunted by up to 50% in response to MDIR (**Fig. 7E and F**). These findings are consistent with our *in vitro* data that demonstrate that 14-3-3ζ is required for accessibility of key adipogenic factors (**Fig. 4-7**). Thus, 14-3-3ζ has a critical role in ensuring the accessibility of key chromatin regions involved in adipogenesis.

## DISCUSSION

Our previous discoveries on the role of 14-3-3ζ in adipogenesis were based on systemic loss and gain of function mouse models (9, 14), and we sought to define which cell type, mature adipocytes or APCs, were key in the regulation of 14-3-3ζ-dependent adiposity. Although deletion of 14-3-3ζ in either cell population affected glucose tolerance, effects on body weight gain and adiposity were primarily seen only when 14-3-3ζ was deleted in APCs. Through the use of CRISPR-Cas9 genome editing, we developed a useful model to explore the nuclear 14-3-3ζ interactome of APCs during adipogenesis, and this led to the discovery that 14-3-3ζ primarily acts during the early stages of adipocyte differentiation to aid in the reorganization of chromatin to allow for adipogenic gene, promoter and enhancer regions to be accessible. These findings, combined with our prior research, broaden our understanding of the importance of 14-3-3ζ in adipocyte development, metabolism, and glucose and lipid homeostasis (9, 11, 15–17). Furthermore, our study clearly demonstrates for the first time that 14- 3-3ζ, a ubiquitously expressed molecular scaffold protein, has critical upstream regulatory roles in adipogenesis.

We first confirmed significant positive correlations between adipose expression of *Ywhaz* and both basal and HF/HS-diet-induced adiposity in mice from the HMDP (26) (**Fig. 1**). This observation aligns with prior measures in metabolically unhealthy human obesity, where expression of 14-3-3ζ and other isoforms were elevated in adipose tissue (60–63). For example, *YWHAZ* was found upregulated, among other upregulated in omental adipose tissue of obese type 2 diabetic patients, as compared with healthy non-obese and non-diabetic obese patients (63). Moreover, a recent GWAS by Ang *et al.* identified *YWHAB* and *YWHAZ* among 74 highly interconnected genes associated with obesity (64). Lastly, elevated expression profiles of 14-3-3 protein-coding transcripts, including *Ywhaz*, have been confirmed in pancreatic islets from human subjects with type 2 diabetes by our and others’ recent studies (13, 65). Whether elevated levels of 14-3-3 transcripts or transcripts variants are causal or associative to adipogenesis and pathogenesis of obesity and type 2 diabetes remains unexplored, but these observations suggest that *YWHAZ* may be a reliable marker for unhealthy metabolic outcomes, such as adiposity, elevated BMI and type 2 diabetes.

Constitutive deletion of 14-3-3ζ in mature adipocytes from male and female mice was without impact on adiposity (**Fig. 2**) and expression of key metabolic genes in adipose tissue (**Sup. Fig. 4**). However, impaired glucose tolerance was detected, suggesting direct or indirect contributions of adipose 14-3-3ζ to whole body glucose uptake. Indeed, previous studies have revealed important contributions of 14-3-3 proteins to the coordination of Rab GTPase-activating protein-dependent pathways that mediate insulin- and exercise-dependent translocation of GLUT transporters for glucose uptake by adipocytes and myocytes (66–70).

Additional studies are required to better assess how adipocyte-specific 14-3-3ζ contributes to whole-body glucose homeostasis, and if it is involved in crosstalk with other metabolically active tissues. For example, both 14-3-3ζ and 14-3-3β have been found in the protein cargos of secreted large adipocyte-derived vesicles (71). Thus, 14-3-3ζ could participate as part of the bioactive adipocyte secretome that modulates glucose and lipid homeostasis, together with adiponectin, leptin, ANGPTL4, and other adipocyte-derived secreted vesicles cargos (72–74). Further supporting the notion that 14-3- 3ζ delivered by EVs could play an important role in metabolism is the finding that 14-3-3ζ was found within human umbilical cord mesenchymal stem cell-derived small extracellular vesicles (hucMSC-sEVs) (75) and that delivery of 14-3-3ζ by EVs diabetic rats with renal dysfunction could improve glycemia, body weight and renal functions (75).

When using a tamoxifen-induced conditional deletion approach, we previously found that 14-3-3ζ deletion in mature adipocytes of adult mice did not result in a notable metabolic phenotype, with the exception of impaired lipolysis (16). In human mature adipocytes, 14-3-3β, ε, γ and ζ were recently identified by Yang *et al.* as interacting proteins with lipid droplets (18). Subsequent functional studies focused on the 14-3-3β isoform and confirmed its requirement for lipolysis (18). Taken together, these findings indicate that in mature adipocytes, 14-3-3 isoforms are primarily involved in adipocyte functions, such as glucose uptake and lipolysis, rather than regulating expansion or proliferation.

In terms of adipose tissue architecture, deletion of 14-3-3ζ in mature adipocytes had no impact when compared with whole-body 14-3-3ζKO mice that were strikingly lean from birth (9). This strongly suggested that 14-3-3ζ was more likely to influence APCs. In fact, we showed that constitutive deletion of 14-3-3ζ in *Pdgfra*+ cells was more impactful on adipocyte size and body weight, but in a sex-specific manner. In males, APC deletion of 14-3-3ζ resulted in reduced body weight and a trend towards lowered fat mass. In female *Pdgfra*14-3-3ζKO mice, fat mass as well as inguinal and perigonadal adipocytes sizes were surprisingly doubled and glucose tolerance markedly impaired. These results reveal that 14-3-3ζ deletion favors adipogenesis in *Pdgfra*+ APCs rather than in mature adipocytes. The sex-specific role of 14-3-3ζ in APCs in white adipose tissue development was surprising, but recent mechanistic studies have revealed regulatory roles of 14-3-3 proteins, including 14-3-3ζ, on the function and localization of estrogen receptor alpha (ERα) and androgen receptor (76, 77). Further studies are required to investigate the contribution of 14-3-3ζ to sex hormones-determined adiposity and sexual dimorphism in adipogenesis. Of note, expression of *Pdgfra* is not limited to APCs but is common to fibroblasts residing in a broad variety of mesenchymal tissues (78). Thus, the *Pdgfra* promoter is not exclusively specific to APCs, and there exists the possibility that the sexual dimorphic effects of deleting 14-3-3ζ in *Pdgfra*+ cells may be mediated by other cell types, aside from APCs.

One possible reason to explain the difference in magnitude or nature of effects between both the *Adipoq*14-3-3ζKO and *Pdgfra*14-3-3ζKO and the whole-body KO mice could be the heterogeneity of the adipocyte and APC populations. In fact, it is now established that heterogenous populations of adipocytes inhabit fat depots, and these adipocytes originate from different APCs (23, 79–81). Even within a given adipocyte (*i.e.*, *Adipoq*+) or APC (*i.e.*, *Pdgfra*+) population, different niches exist with different cellular and physiological fates (23, 79–81). With our previous discovery of heterogenous expression of 14-3-3ζ among mature gonadal adipocytes (16) and the heterogeneous expression of *Ywhaz* among APCs (**Sup. Fig. 3**) (31), this opens new avenue of research to define 14-3-3ζ+ and 14- 3-3ζ- niches and how they contribute to adipocyte development and function.

We further sought to determine the mechanisms by which 14-3-3ζ influences APCs differentiation through deciphering its interactome during the early stages of adipogenesis. Our previous studies have already demonstrated the promising value of mining the 14-3-3ζ interactome to identify novel regulators of adipogenesis (14, 15). For example, we recently identified plakoglobin, the homolog of the adipogenic inhibitor β-catenin, among proteins enriched in the 14-3-3ζ interactome in visceral adipose tissue of mice subjected to HFD (15). To increase the likelihood of detecting novel adipogenic transcriptional regulators, we developed a novel cell model to elucidate the nuclear 14-3-3ζ interactome (**Fig. 4**), and surprisingly, the only transcription factor that could be detected was C/EBP-β (**Sup. Tables 1 and 2**), which confirmed our previous observation (14). Instead, we found that the14-3-3ζ interactome was enriched with regulators of DNA unwinding and chromatin accessibility (**Fig. 4, 6 and 7)**. These findings are consistent with previously reported regulatory functions of 14-3-3 proteins on ribosomal subunits (82) and chromatin modifying enzymes (83–85). Through the use of ATAC-seq, we confirmed the requirement of 14-3-3ζ at early timepoints of adipocyte differentiation in the accessibility and expression of several genes, such as *Fabp4, Adig*, *Retn* and *Fam83a* (**Fig. 6 and 7**). Importantly, all four genes are critical regulators and effectors of adipogenesis.

Our findings pinpoint a new role of 14-3-3ζ in the epigenetic control of adipogenesis (**Fig. 7I**). They support the likelihood that 14-3-3ζ modulates the activity of key chromatin modifying enzymes, including HDAC1 and DNMT proteins (86), that are found in its nuclear interactome. HDAC enzymes consist of 18 mammalian isoforms that promote the condensation of chromatin with discrepant effects on adipogenesis (86). HDAC1 has been shown to inhibit adipogenesis through repressing the C/EBPα promoter (86, 87), while other isoforms, such as HDAC4/5, promote adipogenesis through regulation of GLUT4 promoter (86, 88). The exact mechanisms that control the balance between these isoforms remains poorly defined. Although we did not identify HAT enzymes as part of 14-3-3ζ interactome, they are also of interest as they play antagonist roles to HDAC by promoting chromatin opening. Through targeting H3K9, H3H18 and H3K27 histones, HAT enzymes promote adipogenesis by triggering C/EBPβ and PPARγ expression (86, 89, 90). DNMT enzymes are determinant for lean *versus* obesity-associated adipogenesis (91), and in humans and various preclinical models, DNA hypomethylation of *PPARG*, *ADIPOQ* and *FABP4*, and loss of function of DNMT3A and DNMT1 were reported to provoke aberrant obesogenic growth and impaired metabolic fitness (92–94).

In conclusion, we report novel adipogenic roles of 14-3-3ζ, whereby it primarily functions as a critical regulator of APC fate. Moreover, our findings establish for the first time 14-3-3ζ as a crucial contributor to chromatin remodeling during adipogenesis to allow for the early expression of adipogenic genes. While additional studies are still required to define downstream effectors that mediate 14-3-3ζ’s effects on chromatin remodeling, our proteomic analysis suggest epigenetic factors, such as DNMTs and HDACs, are important in the actions of 14-3-3ζ during APCs differentiation (**Fig. 7G**). Lastly, our study demonstrates the ability of utilizing the 14-3-3ζinteractome to expand our knowledge of processes involved in adipocyte development and, by extension, the pathogenesis of obesity.

## MATERIAL AND METHODS

### Adipose Tissue Gene Expression – Phenotypic correlations in the Hybrid Mouse Diversity Panel (HMDP)

One hundred inbred strains of male and female mice were maintained on a standard lab diet (6% kcal from fat, Ralston Purina Company) until eight weeks of age and subsequently placed on a high-fat /high-sucrose diet(32% kcal from fat and 25% kcal from sucrose) for eight weeks (26). Animals were measured for total body fat mass and lean mass by MRI with Bruker Minispec and software from Eco Medical Systems, Houston, TX every two weeks. After 8 weeks of HF/HS diet, retro-orbital blood was collected under isoflurane anesthesia after 4-hour of fasting. Plasma insulin, glucose, and triglycerides were determined as previously described (95). The HOMA-IR was calculated using the equation [(Glucose × Insulin)/405] (96). Gene expression was measured from flash-frozen perigonadal adipose tissue samples with Affymetrix HT_MG430A. Gene expression-phenotype correlations (biweight midcorrelation) and their significance were calculated with the bicor function in the WGCNA package in R (97).

### Animal husbandry and metabolic testing

*Pdgfra*Cre (C57BL/6-Tg(*Pdgfra*-cre)1Clc/J, strain #013148, Stock #005304, The Jackson Laboratory, Bar Harbor, ME, USA) and *Adipo*qCre (B6.FVB-Tg(*Adipoq*-cre)1Evdr/J, strain #028020, stock #028020, The Jackson Laboratory) mice were bred with Ywhaz^flox/flox^ mice (13, 16) on a C57BL/6J genetic background (Ywhaz^tm1c(EUCOMM)HMgu^; Toronto Centre for Phenogenomics, Toronto, ON, Canada). All mice were maintained on a standard chow diet (Teklad diet no.TD2918) under 12:12-h light-dark cycles in an environmentally controlled setting (21°C) with free access to food and water. All procedures were approved by the Comité institutionnel de protection des animaux (CIPA) du CRCHUM (CIPA protocol #CM20043GLS) and performed in accordance with CIPA guidelines at the University of Montreal Hospital research center.

*Adipoq*Cre:*Ywhaz*^flox/flox^ (*Adipoq*14-3-3ζKO) mice and *Pdgfra*Cre:*Ywhaz*^flox/flox^ (*Pdgfra*14-3-3ζKO) mice, and their respective non-floxed controls (*Adipoq*Cre:*Ywhaz*^wt/wt^ and *Pdgfra*Cre:*Ywhaz*^wt/wt^), were either kept under chow diet until 26 or 16 weeks of age, respectively, weighed weekly, and assessed every 4 weeks for glucose tolerance, insulin sensitivity, and fat and lean mass. For glucose tolerance tests (IpGTT), mice were fasted for 6 h and challenged with D-glucose (2 g/kg b.w.) by intraperitoneal injection. For insulin tolerance tests (IpITT), mice were fasted for 4 h and challenged with Humulin R insulin (0.5 U/kg b.w.; Eli Lilly, Toronto, ON, Canada) by intraperitoneal injection. Blood glucose was measured from the tail vein with a Contour Next EZ glucose meter (Ascencia Diabetes Care, Basel, Switzerland). Body composition was scanned with EchoMRI-100 Body Composition Analyzer (version 2008.01.18, EchoMRI LLC) at the at the Rodent Cardiovascular Core Facility of the CRCHUM. Immediately after euthanasia of mice, perigonadal and inguinal white adipose tissues were collected, weighed, and either flash-frozen in liquid nitrogen for subsequent extraction of RNA or fixed in 4% paraformaldehyde for histological analysis.

### CRISPR-Cas9 knock-in

For the expression of Flag-HA-YWHAZ in 3T3-L1 cell, the method of CRISPR/CAS9 Knock-in was used, Guide RNAs (gRNAs) were designed using the “optimized CRISPR design” tool (https://www.benchling.com/crispr). Oligonucleotides with Bsb1 cleavage overhang were ordered from Life Technologies, annealed, and cloned into Cas9/GFP expressing vector (pX458 from Addgene #48138). The sequence (CATAACTGGATATTCTGTAA) near Start codon ATG were used as gRNA. For the YWHAZ donor template, left homology arms (495bp containing ATG star codon of Ywhaz gene) +Flag-HA + right homology arm (671bp) cassette was synthesized (Bio Basic, Toronto), then cloned into pGEN-HDR-RFP plasmid (dTomato expression plasmid).

The gRNA plasmid (1µg) and donor template plasmid (3µg) were introduced into 3T3-L1 cells (1X 10 6 cell) by electroporation (LONZA, Basel, Switzerland). After 48h of electroporation, GFP and RFP positive cells were sorted into 96 well plates for genotyping and cell culture. Genomic DNA was extracted with QuickExtract (Mandel Scientific, Laval) and PCR was performed with Flag F (caaggacgacgatgacaaagtc) + 3’OUTR (CAGAGCCAACAACGTGAGGT) primers using Q5® High-Fidelity DNA Polymerase (New England Biolabs, MA, USA) according to the manufacturer’s protocol. For Wild type cells, there was no band and Knock in PCR product (992bp) were identified using agarose gel electrophoresis (**Supp fig. 7D**). Genomic DNA was extracted with QuickExtract (Lucigen) and PCR was performed ATG F (TGGAATCTTCTGAACAGGTGGA) + 3’OUTR (CAGAGCCAACAACGTGAGGT) primers using Q5® High-Fidelity DNA Polymerase according to the manufacturer’s protocol. The PCR products were cloned into pMiniT vector. 12 independent recombinant plasmids were selected for Sanger DNA sequencing, with T7 and SP6 used as sequence primers.

### Cell culture, transient transfections, and treatments

3T3-L1, TAP-3T3-L1 and NIH-3T3 cells were kept between passages 10 and 20 for experiments. Cells were maintained in DMEM medium supplemented with 10% newborn calf serum (NBCS) and 1% penicillin-streptomycin. Prior to differentiation, cells were seeded onto 12-well plates or dishes and maintained until confluence. Then, differentiation was induced following the well-established methods of MDI (DMEM supplemented with 10% FBS, 1% P/S, 500 µM IBMX, 500 nM dexamethasone, and 172 nM insulin) or MDIR (DMEM supplemented with 10% FBS, 1% P/S, 500 µM IBMX, 500 nM dexamethasone, and 172 nM insulin, 10µM rosiglitazone) (9, 16, 98, 99). Following MDI or MDIR treatments for 48 hours, the medium was replaced with DMEM supplemented with 10% FBS and 172nM insulin every 2 days. Differentiation was assessed by Oil Red-O staining, as we previously performed, or by measuring expression of key adipogenic genes, 48 hours after MDI or MDIR treatments, by qRT-PCR.

Knockdown of 14-3-3ζ was performed by transfecting cells with siRNA against *Ywhaz* (*siYwhaz,* 25 µM per well, s76189, Silencer Select siRNA, Ambion) or an equivalent concentration of a scrambled control siRNA (#4390844, Silencer Select Negative Control #1, Ambion) using Lipofectamine RNAimax (ThermoFisher Scientific), 24h before differentiation. Overexpression of either *Cebpa, Cebpb, Pparg1* or *Pparg2* in NIH-3T3 cells was performed by transfecting pcDNA3.1(-) rat C/EBP alpha (#12550, Addgene, Watertown, MA, USA), pcDNA-mC/EBPb (#49198, Addgene, Watertown, MA, USA), pSV Sport PPAR gamma 1 (#8886, Addgen, Watertown, MA, USA), or pcDNA3.1-PPARgamma2 (#78768, Addgene, Watertown, MA, USA) plasmids using Lipofectamine 3000 (ThermoFisher Scientific), following manufacturer instructions, 24 hours before transfection of *siYwhaz* and 48 hours before MDIR treatment.

### RNA isolation and qPCR

Total RNA was isolated from 3T3-L1 and NIH-3T3 cells after differentiation and/or transfection or from 10 mg of mice visceral (perigonadal) adipose tissue (VAT), subcutaneous (inguinal) adipose tissue (SAT), or scapular brown adipose tissue (BAT), with Trizol (ThermoFisher Scientific). Afterwards, cDNAs were synthesized with the High-Capacity cDNA Reverse Transcription Kit (ThermoFisher Scientific) prior to qPCR-based measurement of mRNA with SYBR green chemistry on a Quant-Studio 6-flex real-time PCR system (ThermoFisher Scientific). All data were normalized to *Hprt1* or *Actb* by the 2-delta Ct method (100). All primers sequences are listed in supplementary table 3.

### Immunoblotting

TAP-3T3-L1 and 3T3-L1 cells after 0, 24 and 48 hours of MDI and MDIR treatment were lysed followed by a cell fractionation protocol (101). Cytosolic and nuclear protein fractions, supplemented with protease and phosphatase inhibitors, as well as benzonase for the nuclear fraction, (1:50 v:v), were prepared and resolved by SDS-PAGE, transferred to PVDF membranes, and immunoblotted with antibodies against PPARγ1/2, Lamin a/b, and GAPDH. Primary and HRP-linked secondary antibodies were used, and their respective dilutions into blocking solution (I-Block, ThermoFisher Scientific) are listed in supplementary table 4.

### Proteomics (nuclear proteins precipitation, mass spectrometry, and data analysis)

Nuclear extracts were obtained from TAP-3T3-L1 cells (± MDI treatment after 24 or 48h) using extraction buffer which contained 150 mM NaCl, 50 mM Hepes (pH 7.4), 1M Hexylene glycol, 0.5% Sodium deoxycholate, 0.1% SDS, 2 mM EGTA, 1 mM Na_3_VO_4_, and 1 mM N_A_F. A total of amount of 1.5 mg of protein in a 1 mL volume were used for all pull-downs. For each sample, 50 µL of anti-HA magnetic beads (#88836, Pierce) were added, followed by overnight rotation at 4°C. The subsequent day, samples underwent three washes with the extraction buffer and three additional washes with 50 mM NH_4_HCO_3_. Beads were then resuspended in 100 µL of 50 mM NH_4_HCO_3_, to which 1 µg of trypsin was added, followed by overnight incubation at 37°C. An additional 1 µg of trypsin was added the next day, and the incubation continued for 4 hours at 37°C. After centrifugation, the supernatants containing peptides were collected in a separate tube. The beads were then rinsed twice with 100 µL of water, which was added to the collected peptides. Finally, formic acid was added to achieve a final concentration of 4%, and the samples were subjected to speed vacuum concentration prior to analysis by mass spectrometry.

Samples were reconstituted in formic acid 0.2% and loaded and separated on a homemade reversed-phase column (150 μm i.d. x 150 mm) with a 56-min gradient from 0%–40% acetonitrile (0.2% FA) and a 600 nl/min flow rate on an Easy-nLC II (Thermo Fisher Scientific), connected to an Orbitrap Fusion Tribrid mass spectrometer (Thermo Fisher Scientific). Each full MS spectrum acquired with a 60,000 resolution was followed by 20 MS/MS spectra, where the 12 most abundant multiply charged ions were selected for MS/MS sequencing. Raw files were searched with MaxQuant software version 1.3.0.3 (102) and searched against the Uniprot Human database (http://www.uniprot.org/) release 2016_02 (17-Feb-2016) including the reversed database with 74508 entries. General false discovery rate (FDR) for peptides was at 1.0% with decoy removal. Searches were performed at a 25-ppm precursor ion tolerance and 1.0 Da fragment ion tolerance, assuming full tryptic digestion with up to three missed cleavages. Correct peptide identifications were distinguished from incorrect using the target-decoy approach, coupled with linear discriminant analysis as described previously. Once peptides were filtered to an initial FDR of 1%, peptides were assembled into proteins and further filtered to a final protein-level FDR of 1%. All proteins of the same family were grouped, and isoforms of the same protein were considered as one. To estimate the abundance of proteins within eluates, we rely on total spectral counts. Peptides showing significant (*P < 0.05,* non-paired T test) increase in counts at 24h and 48h post differentiation were considered as enriched in 14-3-3ζ interactome at these timepoints. Corresponding protein lists were analyzed with Gene Ontology (ShinyGO, version 0.741) to categorize them on their biological functions.

### Measurements of DNMT, HDAC, or HAT activity

Nuclear protein fractions were isolated from 3T3-L1 cells, plated on 100 mm diameter dishes, treated with scrambled siRNA or si14-3-3ζ, at 0, 24 and 48 hours of MDI-induced differentiation. Equal quantities of proteins were subjected to a DNA methyltransferase (DNMT) colorimetric activity assay (#ab113467Abcam, Toronto, ON, Canada,), Histone deacetylase (HDAC) fluorometric activity assay (#ab156064, Abcam Toronto, ON, Canada) and Histone acetyltransferase (HAT) colorimetric activity assay (#ab65352, Abcam Toronto, ON, Canada)

### ATAC-seq and RNA-seq

Assay for Transposase-Accessible Chromatin using high-throughput sequencing (ATAC-seq) was performed following protocols adapted from Buenrostro *et al*. (103), and Batie *et al* (104). Following treatments, 3T3-L1 cells were washed on plates with PBS and exposed to cell lysis buffer (10 mM Tris-HCl, pH 7.4, 10 mM NaCl, 3 mM MgCl2,0.1% IGEPAL CA-630). After 10 minutes of incubation on ice nuclei were detached, centrifuged (600g, 10 min, 4℃), and washed twice in PBS with repeated centrifugation. Pellets (nuclei fractions) were resuspended in transposition buffer (50% v/v 2X Tagment DNA (TD) Buffer (Illumina, 336 Cambridge, UK), 32% v/v PBS, 0.1% v/v Tween-20, 0.1 mg/mL Digitonin (Promega, Southampton, UK), supplemented with 5% v/v TDE1 Tagment DNA Enzyme (Illumina, Cambridge, UK) in nuclease free water (Sigma, Gillingham, UK)), by gentle pipetting. The transposition reaction was initiated by incubation at 37℃ and stopped by cooling to 4℃. Following DNA extraction, tagmented DNA was amplified by PCR in a reaction mix (5 µL DNA, 2.5 µL of 25 μM forward primer (Nextera/Illumina i5 adaptors (Illumina)), 2.5 µL of 25 μM reverse primer (Nextera/Illumina i7 adaptors (Illumina), 10 µL of SYBR green reagent) with the following thermocycler program: 5min 72°C, 30sec 98°C and 11 cycles of 10sec 98°C, 30sec 63°C and 1min 72°C.

Adaptors were trimmed from the ATAC-seq reads using Trim Galore (https://github.com/FelixKrueger/TrimGalore) and reads aligned to mm10 with Bowtie2 (105). Picard (http://broadinstitute.github.io/picard/) was used to assess library size distribution. Bowtie2 was also used for the alignment of 3T3-L1 adipocytes ChIP-seq reads: GSM535755 (H3K4me3) GSM535758 (H3K27ac) and GSM535740 (input) downloaded from series GSE21365 (106). MACS2 (107) was used for peak calling. Quantification of peaks-associated ATAC-seq reads was performed using featureCounts (108). RNA-seq data from GSE60745 (9) was realigned and quantified using Salmon (109). Differential analysis of quantified peaks (ATAC-seq) and gene expression (RNA-seq) was performed with DESeq2 (110). Bedtools (111) was used to integrate ATAC-seq and ChIP-seq. Transcript factor motif analysis and gene annotation of peaks for integration with gene expression data was performed with the annotatePeaks function of HOMER (112). Correlation analysis was performed in GraphPad prism with all annotated peaks or with a subset of peaks annotated within 1500 nucleotides of the closest transcription start site (TSS). Differentially open and closed regions were submitted to ShinyGO v0.741 for KEGG pathway enrichment characterization.

### Single nuclei RNA-seq analysis to determine *Ywhaz* heterogeneity in APCs

Filtered datasets including metadata consisting of mature adipocytes or APC populations from 20-week-old mice of both sexes were obtained from Emont *et al*. (31), and further sub-setted to contain only chow fed animals. Samples raw counts were then extracted and normalized using SCTransform method (113) and integrated with Seurat v5 (114). Data were visualized using Uniform Manifold Approximation and Projection (UMAP) (115) and the normalized and log-transformed expression values were used for downstream visualizations.

## Data availability

Proteomics data were deposited in ProteomeXchange through partner MassIVE as a complete submission and assigned MSV000094119 and PXD049695. It can be downloaded from ftp://MSV000094119@massive.ucsd.edu, using the password: Rial.

Assay for Transposase-Accessible Chromatin using high-throughput sequencing (ATAC-seq) data were deposited to the Gene Expression Omnibus.

## Statistical Analysis

All data are expressed as the mean ± s.e.m. Data were analysed by ANOVA followed by Dunnett or Bonferroni t-tests, or by Student’st-tests, and significance was achieved when P ≤ 0.05. A minimum of n= 4 independent experiments was performed for analysis.

## Acknowledgements

This work was supported by CIHR Project (PJT-153144) and NSERC Discovery (RPGIN-2017- 05209) to GEL, as well as funding from the Cardiometabolic Health Diabetes, and Obesity (CMDO) Reserch Network (Jean-Davignon Young Investigator Award) and the Banting Research Foundation (Discovery Award). Infrastructure support was provided through a Canada Foundation for Innovation (CFI)-John Evans Leaders Fund (JELF-37405) to GEL. GEL holds the Canada Research Chair in Adipocyte Development. SAR was previously supported by a postdoctoral award from the Fonds de Recherche du Québec Nature et technologies (FRQNT). AV was supported by a Mitacs Accelerate Postdoctoral Award and by the Banting Research Foundation. The authors acknowledge Drs. Roger Cox and Samantha Laber (MRC Harwell Institute, University of Oxford, UK) for discussions with ATAC-seq and sharing an optimized experimental protocol. The authors also acknowledge Dr Nhung Vuong- Robillard (CRCHUM) who developed and optimized the macros for morphological analysis of adipose tissue histosections with Visomorph™. MC was supported by NIH grant R01DK118287 as well as American Diabetes Association Innovative Basic Science Award (1-19-IBS-105). PPR is supported by a CIHR Project grant (PJT-178154). AM-S was supported by a New Investigator Research Grant (MR/P023223/1) and a Project Grant (MR/X009912/1) by the Medical Research Council (MRC, UK). Proteomic analyses were performed by the Center for Advanced Proteomics Analyses (CAPA), a Node of the Canadian Genomic Innovation Network that is supported by the Canadian Government through Genome Canada.

## CONTRIBUTIONS

SAR and GEL designed the study, collected, and organized the results, and wrote the manuscript. SAR, ZY, VA, DS, AA, GL, MC, AM-S, PPR, TMD, GEL performed experiments, analyzed data, and/or edited the manuscript. GEL is the guarantor of this work.

---

**Supplementary figure 1:**
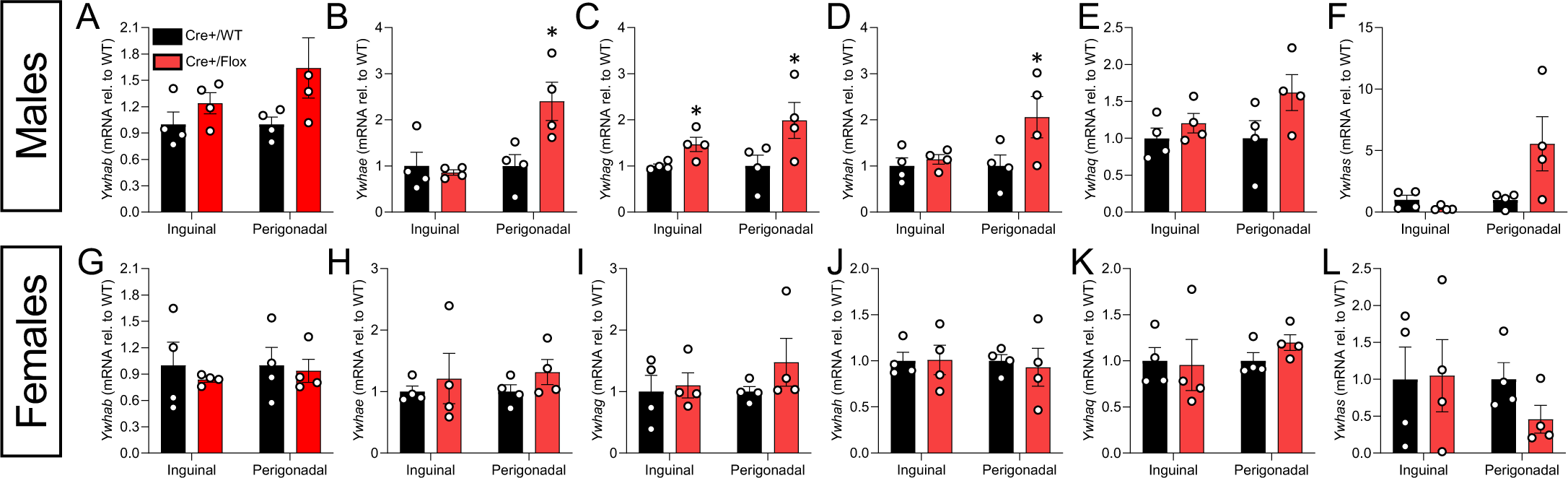
Impact of 14-3-3ζ deletion in mature adipocytes on the expression of remaining 14-3-3 isoforms mice. **(A-F)** Male and **(G-L)** female wild type (*Adipoq*-Cre+/WT) and *Adipoq14-3-3*ζKO (*Adipoq*-Cre+/Flox) mice were fed a chow diet until 26 weeks of age. Inguinal (SAT) and perigonadal (VAT) adipose tissue samples were subjected to qRT-PCR to measure *Ywhab, Ywhae, Ywhazg, Ywhah, Ywhaq and Ywhas* mRNA levels following the delta(Δ) CT method with murine *Hprt1* as the housekeeping gene (n=4 per sex per genotype). Error bars represent S.E.M. Significant differences between wild type and Adipoq*14-3-3*ζKO mice are indicated by \**P*<0.05 (calculated by Student’s T-test).

**Supplementary figure 2:**
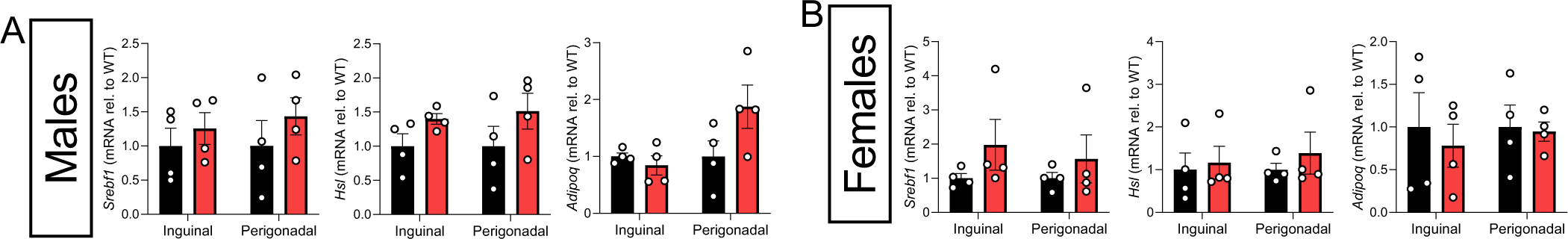
Deletion of 14-3-3ζ in mature adipocytes does not modulate the expression of key genes of adipose homeostasis. Inguinal (SAT) and perigonadal (VAT) adipose tissue samples extracted from 26-weeks old **(A)** male and **(B)** female wild type (*Adipoq*-Cre+/WT) and *Adipoq14-3-3*ζKO (*Adipoq*-Cre+/Flox) mice were subjected to qRT-PCR to measure *Srebf1, Hsl and Adipoq* mRNA levels following the delta(Δ) CT method with murine *Hprt1* as the housekeeping gene (n=4 per sex per genotype). Error bars represent S.E.M

**Supplementary figure 3:**
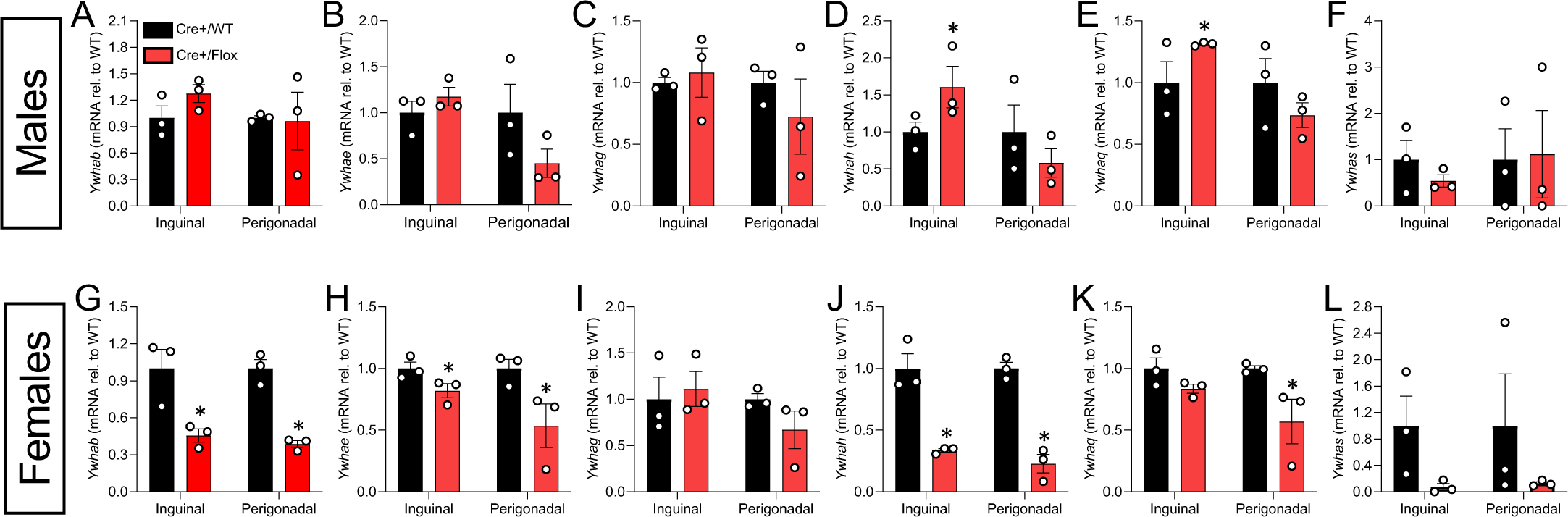
Impact of 14-3-3ζ deletion APC on the expression of remaining 14-3-3 isoforms mice. **(A-F)** Male and **(G-L)** female wild type (*Pdgfra*-Cre+/WT) and *Pdgfra14-3-3*ζKO (*Pdgfra*-Cre+/Flox) mice were fed a CHW diet until 26 weeks of age. Inguinal (SAT) and perigonadal (VAT) adipose tissue samples were subjected to qRT-PCR to measure *Ywhab, Ywhae, Ywhazg, Ywhah, Ywhaq and Ywhas* mRNA levels following the delta(Δ) CT method with murine *Hprt1* as the housekeeping gene (n=3-4 per sex per genotype). Error bars represent S.E.M. Significant differences between wild type and *Pdgfra*14-3-3ζKO are indicated by \**P*<0.05 (calculated by Student’s T-test). Error bars represent S.E.M.

**Supplementary figure 4:**
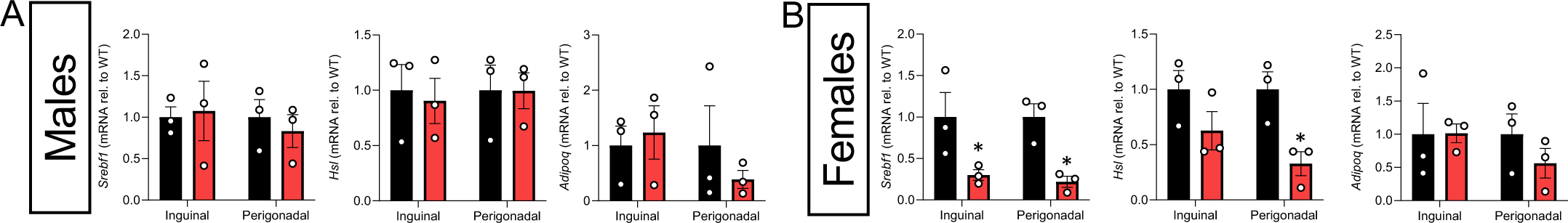
Deletion of 14-3-3ζ in adipocyte progenitor impacts the expression of adipose key metabolic genes in a sex-specific manner. Inguinal (SAT) and perigonadal (VAT) adipose tissue samples extracted from 26-weeks old **(A)** male and **(B)** female wild type (*Pdgfra*- Cre+/WT) and *Pdgfra14-3-3*ζKO (*Pdgfra*-Cre+/Flox) mice were subjected to qRT-PCR to measure *Srebf1, Hsl and Adipoq* mRNA levels following the delta(Δ) CT method with murine *Hprt1* as the housekeeping gene (n=4 per sex per genotype). Error bars represent S.E.M. Significant differences between wild type and *Pdgfra*14-3-3ζKO are indicated by \**P*<0.05 (calculated by Student’s T-test).

**Supplementary figure 5:**
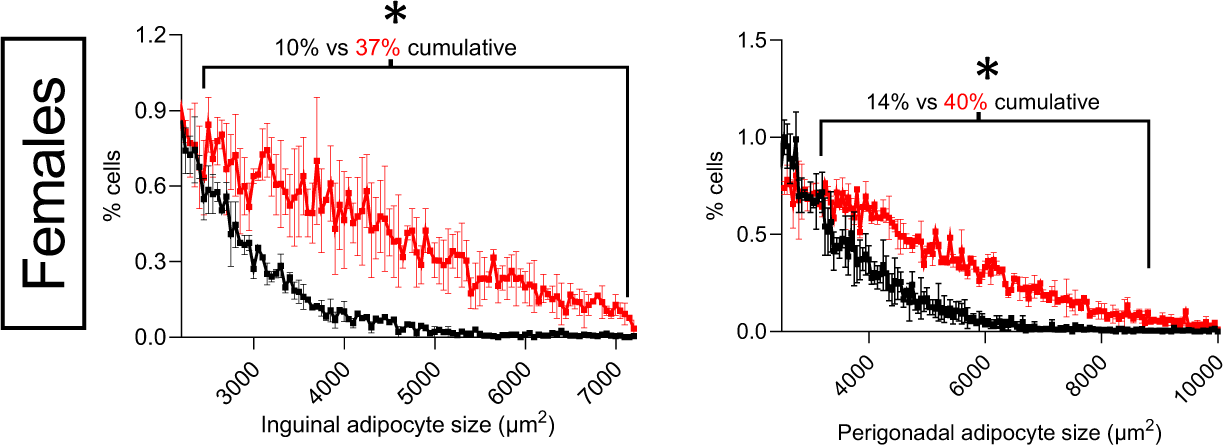
14-3-3ζ deletion in adipocyte progenitor cells results in adipocytes hypertrophy in female mice. Size distribution of large (≥ 2,500 µm^2^) inguinal (SAT) and perigonadal (VAT) adipocytes in histological sections of adipose depots extracted from 26-weeks old female *Pdgfra*- Cre+/WT and *Pdgfra*-Cre+/Flox mice.

**Supplementary figure 6:**
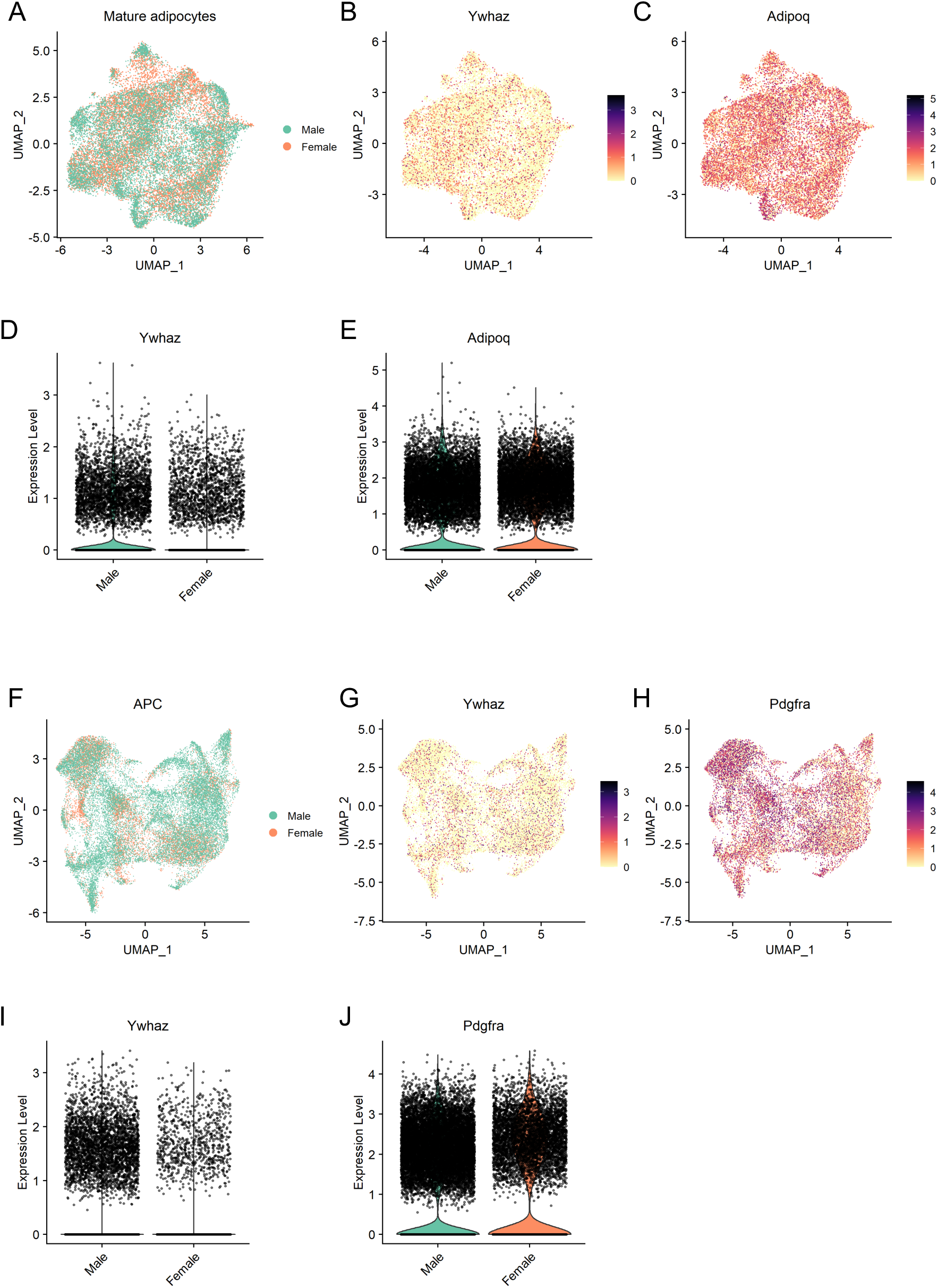
14-3-3 is heterogeneously expressed within mature adipocytes and APC clusters of male and female males. Single-nucleus transcriptomic analysis of mouse mature adipocytes and APCs. (**A**) UMAP plot of pooled male and female populations of mature adipocytes. (**B-C**) UMAP plot of pooled male and female mature adipocytes showing expression of *Ywhaz* and *Pdgfra.* (**D-E**) Violin plots showing the expression of *Ywhaz* and *Pdgfra* in male and female mature adipocytes separately. (**F**) UMAP plot of pooled male and female APC populations. (**G-H**) UMAP plot of pooled male and female APCs showing expression of *Ywhaz* and *Pdgfra*. (**I-J**) Violin plots showing the expression of *Ywhaz* and *Pdgfra* in male and female APCs separately.

**Supplementary figure 7:**
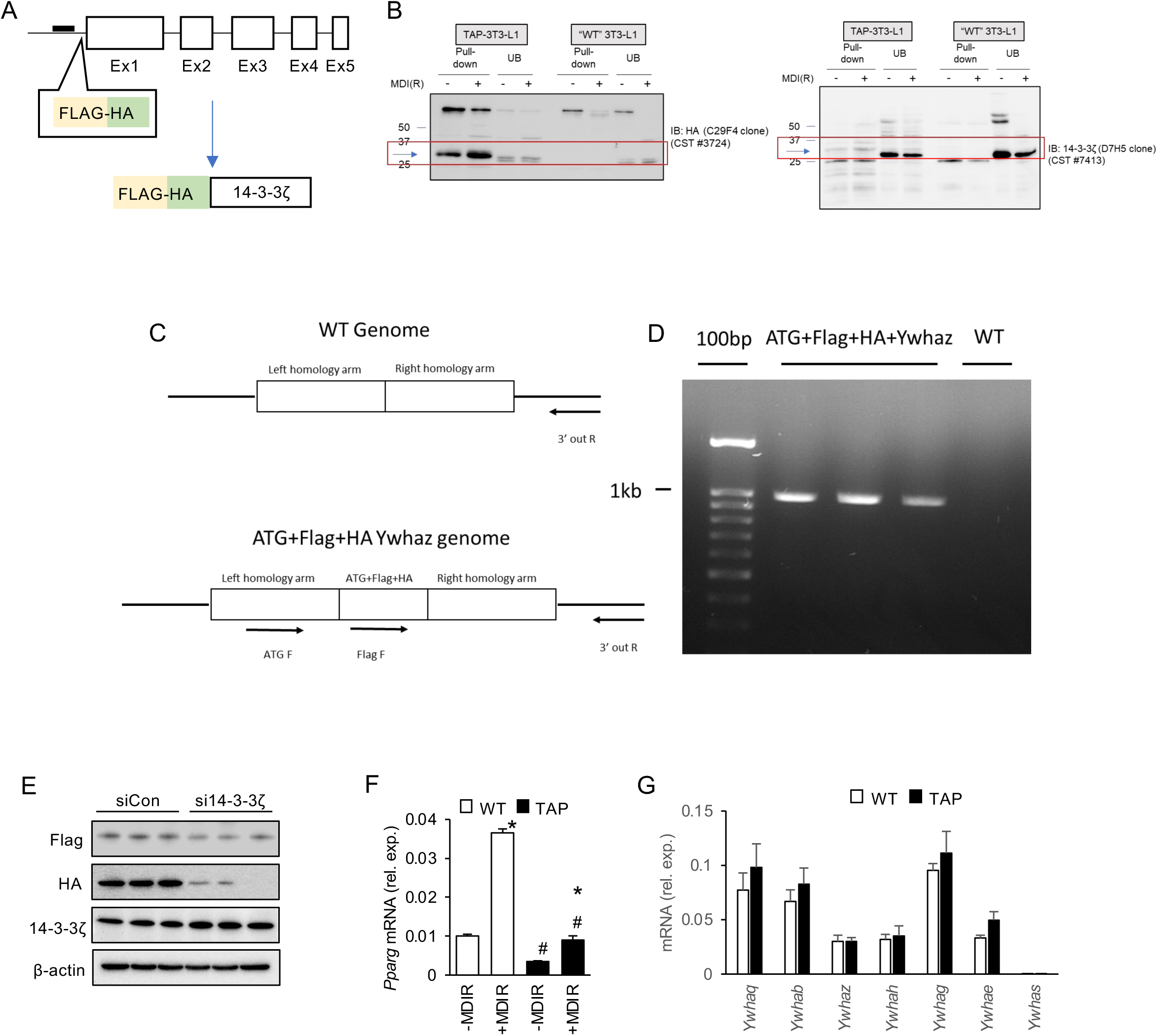
Validation of CRISPR-Cas9-mediated insertion of a TAP (FLAG-HA) tag at the locus of *Ywhaz* in 3T3-L1 cells. (**A)** Schematic representation of the CRISPR-Cas9-mediated knock-in of TAP(FLAG-HA)-coding sequence in 5’ of exon1 of Ywhaz in TAP-3T3-L1 cells. (**B**) validation by anti-HA pull down followed by Western blot of expression of a TAP-tagged 14-3-3ζ by TAP-3T3-L1 cells. (**C and D**) Confirmation by PCR of the knock-in of TAP-coding sequence downstream the ATG initiation codon of *Ywhaz*. (**E**) Concomitant loss of 14-3-3ζ protein and TAP subunits (FLAG and HA) in TAP-3T3-L1 cells treated with *siYwhaz,* assessed by Western blot. (**F**) Validation by qRT-PCR of conserved MDIR-dependent induction of *Pparg* gene expression in TAP-3T3-L1 cells as compared with non-transformed native 3T3-L1 cells. (G) Validation by qRT-PCR of normal stoichiometry of *Ywha* genes isoforms in TAP-3T3-L1 cells. Error bars represent S.E.M. (**F**) Significant differences between MDIR- treated and non-treated cells are indicated by \**P*<0.05 and significant differences WT- and TAP-3T3-L1 cells are indicated by ^#^*P*<0.05 (calculated by Student’s T-test) (n=3 per group).

**Supplementary figure 8:**
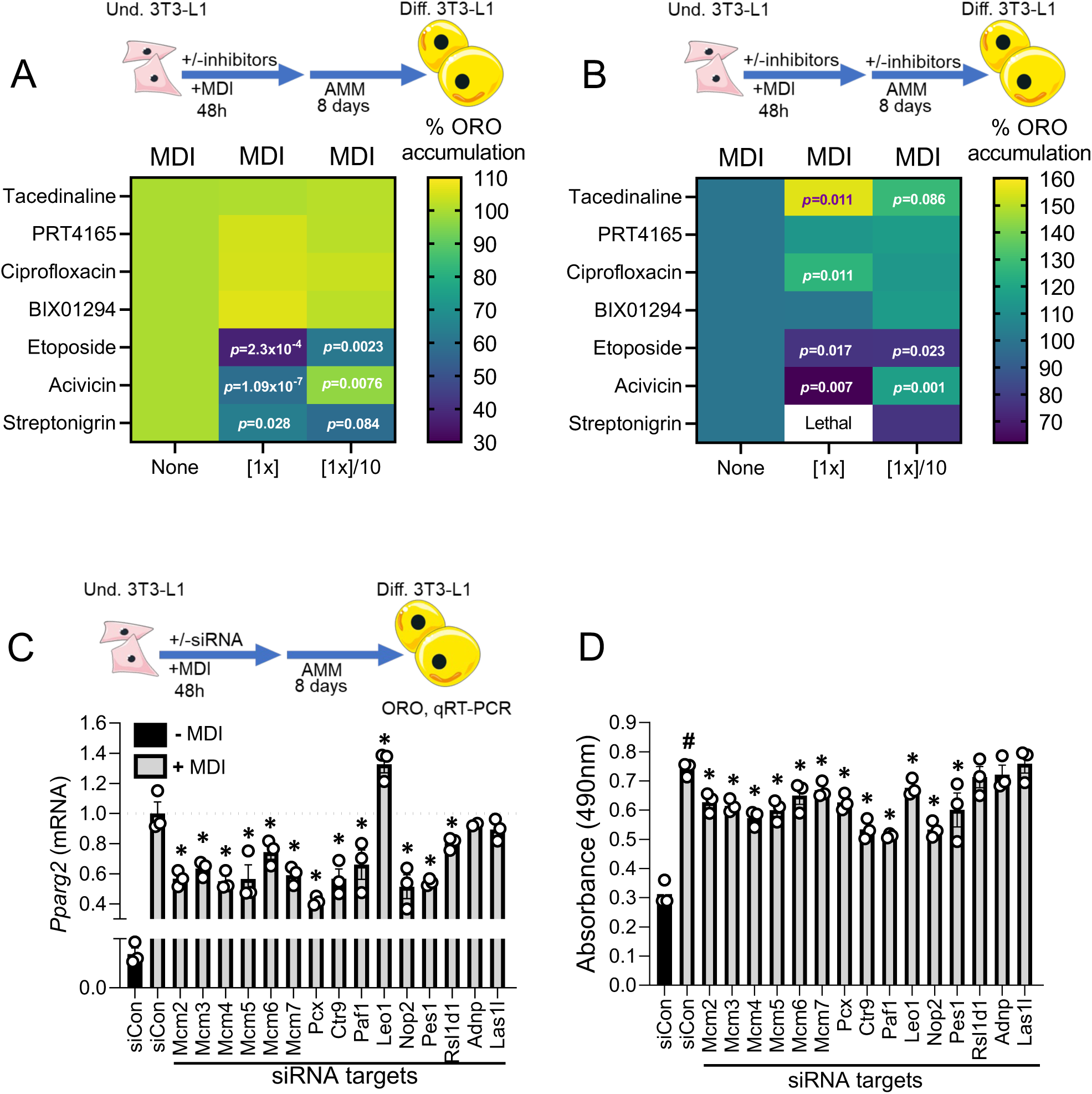
DNA unwinding, histone deacetylase activity, and ribosomal assembly, annotated as biological processes of the 14-3-3ζ nuclear interactome, are important modulators of adipogenesis. (**A,B**) Heat maps showing the percentage of Oil Red O incorporation by 3T3-L1 cells treated with MDI relative to non-treated cells. MDI-treated cells were exposed to 7 compounds at inhibiting concentrations (on respective biological functions) during the first 48h (**A**) or the 10 days (**B**) of MDI-treatment protocol. Doses of the 7 compounds corresponding to respective IC_50_ are identified as [1X] and doses corresponding to IC_50_/10 are identified as [1X]/10 (**A,B**, n=3 biological replicates per condition). (**C,D**) *Pparg2* gene expression measured by RT-qPCR (C) and Oil Red O incorporation (D) levels measured by absorbance at 490nm, in MDI-differentiated 3T3-L1 cells exposed to either gene-targeting siRNA (25 nM) or equal concentration of siCTL during the first 48h of differentiation (n=3 biological replicates per condition). Significant differences between MDI-differentiated cells treated with gene-targeting siRNA and MDI-differentiated cells treated with siCTL are indicated by \**P*<0.05. Significant differences between siCTL and MDI-treated cells and siCTL non-MDI-treated cells are indicated by ^#^*P*<0.05 (calculated by Student’s T-test, n=3 biological replicates per condition).

